# Full-length transcriptomic analysis in murine and human heart reveals diversity of PGC-1α promoters and isoforms regulated distinctly in myocardial ischemia and obesity

**DOI:** 10.1101/2022.03.23.485511

**Authors:** Daniel Oehler, André Spychala, Axel Gödecke, Alexander Lang, Norbert Gerdes, Jorge Ruas, Malte Kelm, Julia Szendroedi, Ralf Westenfeld

**Affiliations:** Division of Cardiology, Pulmonology, and Vascular Medicine, Medical Faculty, Heinrich-Heine University, Düsseldorf, Germany; Cardiovascular Research Institute Düsseldorf (CARID), Medical Faculty, Heinrich-Heine University, Düsseldorf, Germany; Department of Cardiovascular Physiology, Heinrich-Heine University Düsseldorf, Düsseldorf, Germany; Molecular and Cellular Exercise Physiology, Department of Physiology and Pharmacology, Karolinska Institutet, SE-17177, Stockholm, Sweden; Institute for Clinical Diabetology, German Diabetes Center, Leibniz Center for Diabetes Research, Heinrich Heine University, Düsseldorf, Germany

**Keywords:** Ischemia/Reperfusion, Long-Read Sequencing, PGC-1α, Diet-Induced Obesity

## Abstract

**Background:** Peroxisome proliferator-activated receptor gamma coactivator-1 alpha (PGC-1α) acts as a transcriptional coactivator and regulates mitochondrial function. Various isoforms are generated by alternative splicing and differentially regulated promoters. In the heart, total PGC-1α deficiency knockout leads to dilatative cardiomyopathy, but knowledge on the complexity of cardiac isoform expression of PGC-1α remains sparse. Thus, this study aims to generate a reliable dataset on cardiac isoform expression pattern by long-read mRNA sequencing, followed by investigation of differential regulation of PGC-1α isoforms under metabolic and ischemic stress, using high-fat-high-sucrose-diet-induced obesity and a murine model of myocardial infarction.

**Methods and Results:** Murine (C57Bl/6J) or human heart tissue (obtained during LVAD-surgery), was used for long-read mRNA sequencing, resulting in full-length transcriptomes including 58,000 mRNA isoforms with 99% sequence accuracy. Automatic bioinformatic analysis as well as manual similarity search against exonic sequences lead to identification of putative coding PGC-1α isoforms, validated by PCR and Sanger-Sequencing. Thereby, 12 novel transcripts generated by hitherto unknown splicing events were detected. In addition, we postulate a novel promoter with homologous and strongly-conserved sequence in human heart. High-fat-diet as well as ischemia/reperfusion (I/R) injury transiently reduced cardiac expression of PGC-1α-isoforms, with the most pronounced effect in the infarcted area. Recovery of PGC-1α-isoform expression was even more decelerated when I/R was performed in diet-induced obese mice.

**Conclusions:** We deciphered for the first time a complete full-length-transcriptome of the murine and human heart, identifying novel putative PGC-1α coding transcripts including a novel promoter. These transcripts are differentially regulated in I/R and obesity suggesting transcriptional regulation and alternative splicing that may modulate PGC-1α function in the injured and metabolically challenged heart.

## Introduction

The ubiquitously expressed Peroxisome proliferator-activated receptor gamma coactivator-1 alpha (PGC-1α) regulates mitochondrial function, hypoxia-induced angiogenesis and antioxidative capacity. It functions as a transcriptional coactivator and its transcripts are assembled by alternative splicing events and are regulated by different promoters(1–3). Its biological roles range mitochondrial biogenesis and mitochondrial dynamics(4) through induction of hypertrophy and atrophy in muscle cells as well as controlling inflammation in apoptosis environments(5). PGC-1α is studied in detail in liver, skeletal muscle, neuronal cells (neurodegenerative disorders)(6, 7) and cancer(8).

In skeletal muscle in mice, general activation of PGC-1α leads to protection from sarcopenia(9). Additionally, PGC-1α activation in muscle promotes angiogenesis, oxygen consumption and energy supply(10–12). Moreover, it leads to an anti-inflammatory environment(13), while loss of whole PGC-1α in muscle potentiates a systemic inflammatory response(14).The most intensively studied isoform, PGC-1α1 (also known as PGC-1α-a), leads in cultured muscle cells and in skeletal muscle *in vivo* to an increased expression of several angiogenic factors (including VEGF), and consecutively to accelerated recovery after ischemia, which is severely impaired in PGC-1α-deficient mice(15, 16). Interestingly, the short isoform PGC-1α4 induces muscle hypertrophy, most likely via a negative feedback mechanism to myostatin expression(1). Thus, in skeletal muscle, differential usage of isoforms generated through alternative splicing have opposite effects, depending on the expressed isoform. Besides alternative splicing, differential promoter usage represents another level of regulation, as for example systemic cold stress leads to increased expression of PGC-1α isoforms controlled by the alternative promotor(17).

The heart is crucially depending on energy supply, mitochondrial oxidative capacity, and mitochondrial biogenesis. The PGC-1 family of transcription factors is involved in cardiac metabolism as a main driver of mitochondrial biogenesis and induces scavengers of ROS-species(18, 19). Additionally, it is involved in indirect transcriptional regulation of the mitochondrial genome, import and utilization of fatty acids and angiogenesis(20). Moreover, previous work showed a relation between deficiency of total PGC-1α and impaired mitochondrial function(21–23). Transgenic mice with myocardial inactivation/deletion/deficiency of total PGC-1α develop a dilatative cardiomyopathy phenotype with increased end systolic volume and reduced contractile function as well as metabolic alterations associated with heart failure(24), while overexpression of PGC-1α-a appears to enhance contractility without negative feedback on cardiac metabolism(25).

The crosstalk between metabolic dysfunction and heart failure in terms of ‘diabetic cardiomyopathy’ is still not fully understood(26). PGC-1α hereby mediates adaptation to caloric restriction(27) and suppresses inflammatory processes mediated by NFκB(28) in mice fed with high-fat diet. Additionally, PGC-1α has a protective effect on mitochondrial function in insulin resistance(29). Considering that in skeletal muscle alternative splicing and varying promoters generate different isoforms with distinct functions, we aimed to investigate the detailed regulation of PGC-1α expression (and function) also in heart.

As neither in-depth information about the cardiac transcriptome in general nor data on possible differential expression for PGC-1α in particular were available, this study is the first to fill this gap by latest-generation sequencing using long-read full-length transcriptomics. Second, as the functional impact of differential-expressed isoforms PGC-1α in the heart remains elusive, we investigate modified expression patterns under metabolic and cardiovascular challenges in cardiac tissue. Using ischemia/reperfusion (I/R) injury alone or in combination with diet-induced obesity and pre-diabetes we aim model pathological conditions such as acute myocardial infarction in diabetes-prone patients.

## Results

### First full-Length Transcriptome in murine heart reveals over 58000 unique isoforms

Classical short-read sequencing techniques are not suited to generate full-length transcriptomes with reliable sequence information, which is necessary for detection of diverse splice variants. Thus, high accuracy on sequence level is crucial for downstream experiments. We generated a comprehensive full-length transcriptome map using mRNA long read sequencing (Figure 1, A). In murine wildtype heart, we identified 58440 unique isoforms with 99 % predicted accuracy, originating from 12789 genes (Figure 2, A-C). Exploring the underlying mechanism of transcript formation, we identified 117182 known canonical (92.04 %), 57 known non−canonical (0.04%), 6306 novel canonical (4.95%) and 3768 novel non−canonical (2.96%) splice junctions.

**Figure 1:**
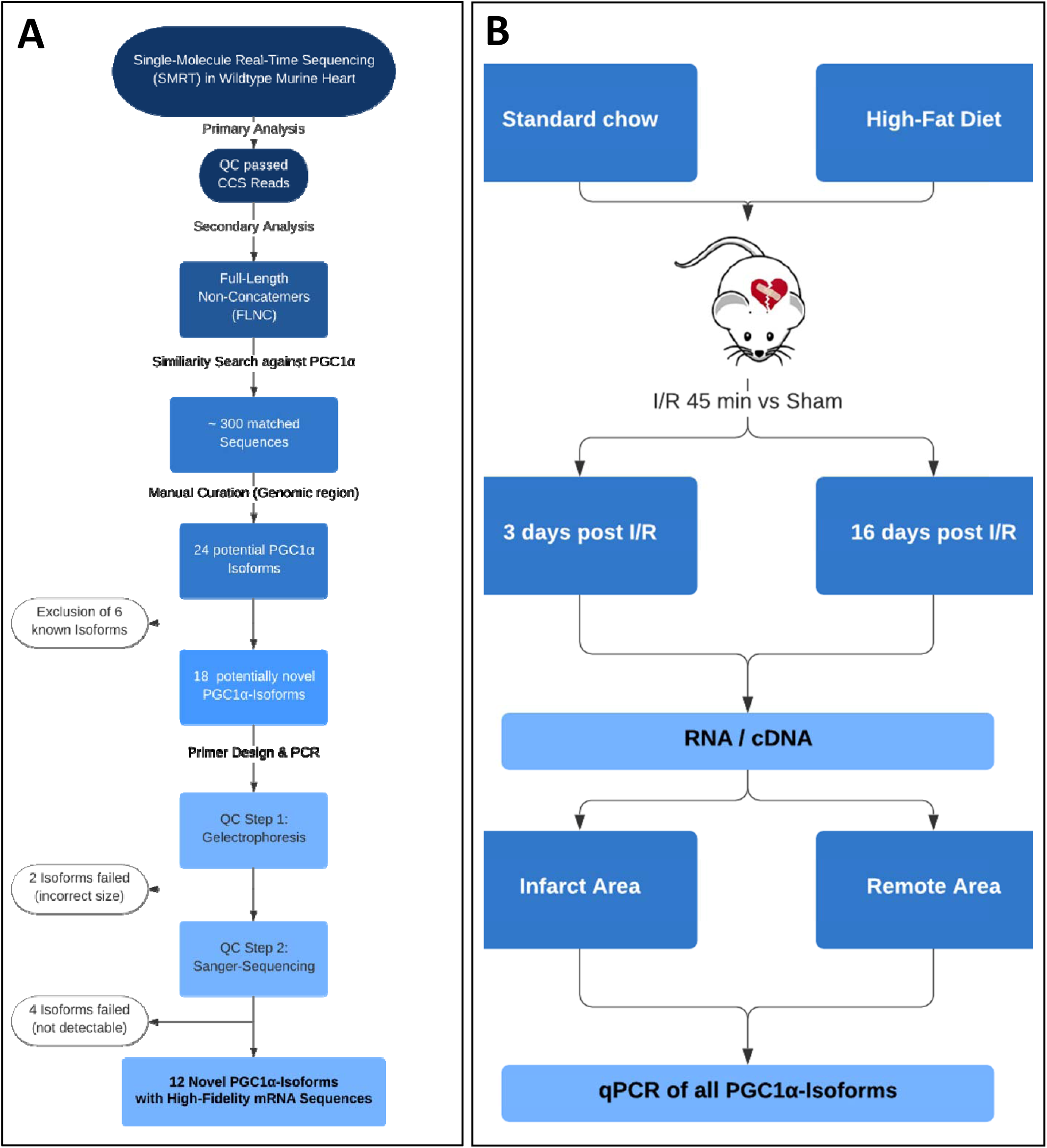
Workflow overview. **A.** Strategy for detection and validation of PGC-1α -isoforms by SMRT-Sequencing in mice. Starting from raw data from SMRT-Sequencing, primary and secondary analysis were performed, resulting in Full-Length Non-Concatemers (FLNC). Then, a similarity search against PGC-1α was performed, yielding in 18 potential novel PGC-1α transcripts, resulting in 12 high-fidelity novel PGC-1α isoforms after quality control. **B.** Strategy for investigating into differential expression of PGC-1α -isoforms by diet-induced obesity and Ischemia/Reperfusion (I/R) injury in mice followed by qPCR. Mice fed either a standard chow or high-fat diet underwent I/R or sham-surgery. Then, tissue from the infarct area and the remote area (distant from the infarcted area) as baseline as well as 3 and 16 days post I/R was collected and used for expression analysis using qPCR.

**Figure 2:**
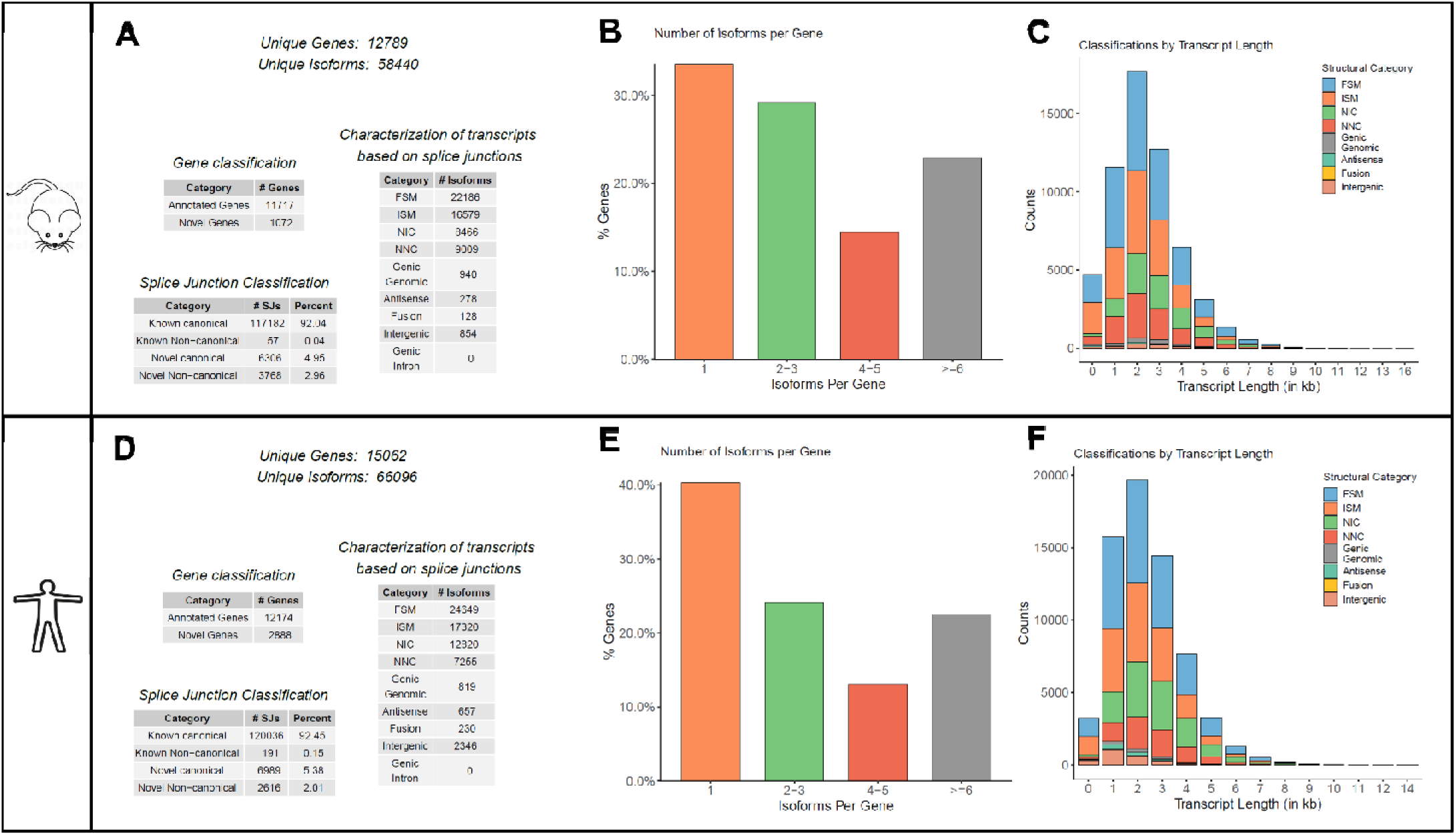
Analysis of SMRT-Sequencing in murine and human heart. Unique genes and isoforms and their characterization from automated analysis of murine (**A-C**) and human (**D-F**) datasets. Shown is the numeric classification of found genes and isoforms (**A** and **D**), number of isoforms per gene (**B** and **E**) and length-distribution of transcripts (**C** and **F**). Details see text.

The majority of all found isoforms (n= 22186) used junctions and corresponding exons matching the annotation (‘Full-Splice-Match”, FSM). Respectively, 16579 isoforms used known splice junctions in consecutive order with some parts missing (e.g. last part of a transcript; ‘Incomplete Splice Match”, ISM). The minority of all isoforms were either novel in catalog (NIC, n=8466), using known splice junctions, but resulting in different transcripts, or novel not in catalog (NNC, n=9009), using new splice junctions and resulting in previously not annotated transcripts. The overall quality of the dataset was excellent (see Supplemental Figure S01).

### Deciphering novel Full-Length Transcriptome in human heart by Long-Read sequencing

As knowledge on human data is even more limited, we aimed to unravel the human heart transcriptome by means of the same approach. Thus, we created a transcriptomic map using mRNA derived from left ventricle from one patient undergoing LVAD-Implantation. In human heart we identified 66096 unique isoforms originating from 15062 genes (Figure 2, D-F). Here, 120036 known canonical (92.45%), 191 known non−canonical (0.15%), 6989 novel canonical (5.38) and 2616 novel non−canonical (2.01%) splice junctions were detected. The majority of all found isoforms (n= 24649) was categorized as FSM, 17320 isoforms as ISM. A substantial part of all isoforms was either NIC (n=12820) or NNC (n=7255). The overall quality of the transcriptomic data was also very good and comparable to the murine dataset (Supplemental Figure S01).

### Detection of 12 novel PGC-1α-Isoforms within the Full-Length-Transcriptome of the Murine Heart

Within our novel and complete cardiac transcriptome, we further evaluated expression of PGC-1α as an example for a complex genomic region/structure yielding highly diverse isoforms. These isoforms derive from multiple splice events as well as different promoter sites and regulate diverse biological functions in vivo(1–3). Two approaches were performed (see also workflow in Figure 1, A): First, we used our high-quality, full-length, clustered transcripts after mapping to reference genome and filtered for those with potential Open-reading Frame (ORF). Due to a potential loss of isoforms during filtering steps of the automated pipeline we further performed a similarity search against PGC-1α within reads without previous clustering and mapping (full-length, non-concatemers; FLNC). As these reads are non-polished and therefore not error-corrected, manual curation was necessary. This led not only to the identification of six known but also remarkably 18 potentially novel PGC-1α transcript variants with valid ORF prediction (Figure 1, A). We confirmed cardiac expression of 12 of the predicted novel transcripts using qPCR. (Figure 3 and supplemental Figure S02).

**Figure 3:**
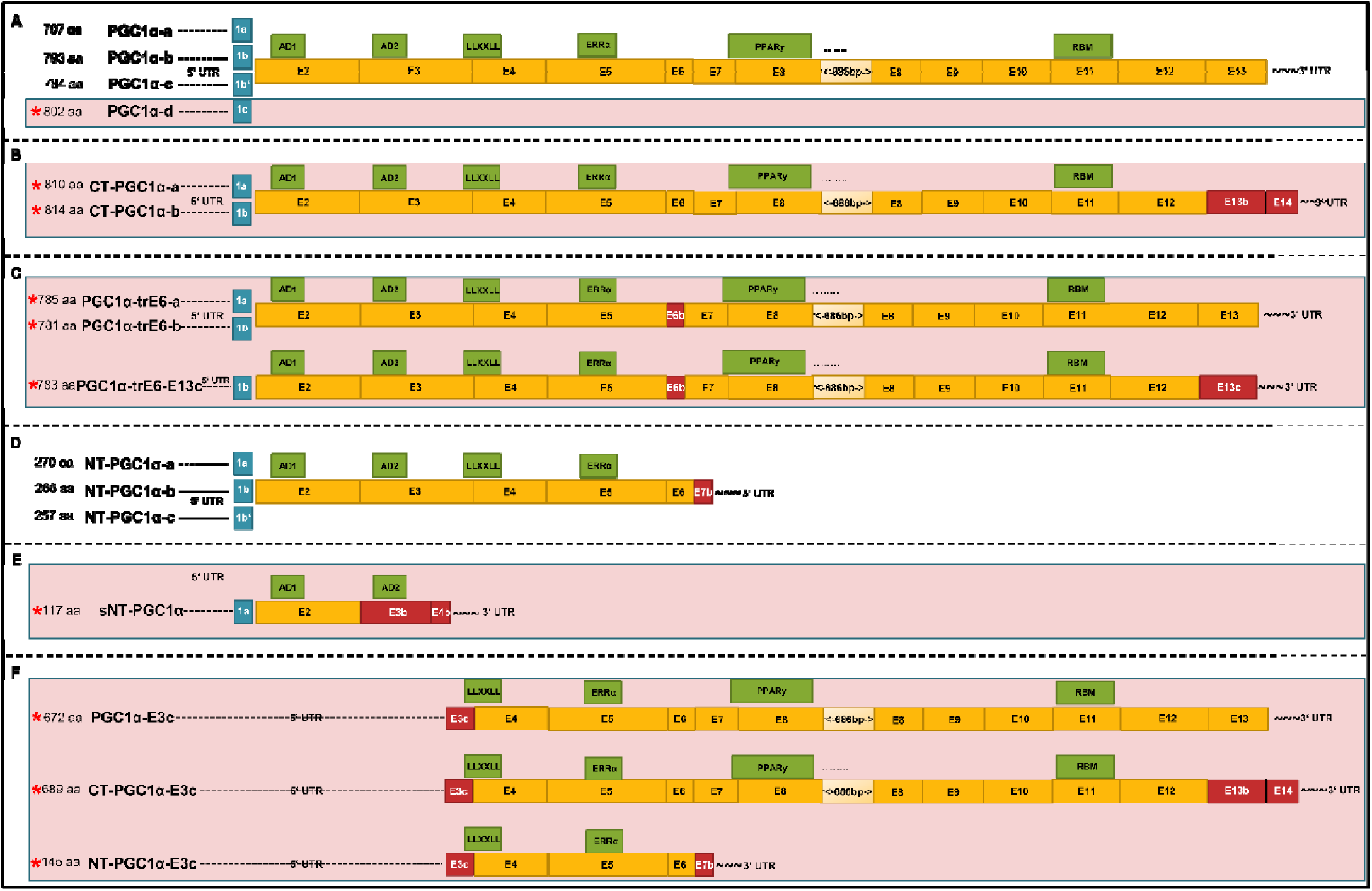
Overview over PGC-1α-Isoforms in murine heart (passed QC) PGC-1α Isoforms (mRNA) after SMRT-Sequencing which passed QC-Filtering (Figure 1): Differing starting exons 1a, b, b’ or c (blue boxes), canonical main exons (orange boxes), novel/altered exons (red boxes) and functional domains (green boxes, details see text). Length of boxes indicates relative length of nucleotides (true to scale, with exception of exon 8: shortened bp marked), asterisks and red background layer indicating novel isoforms. **A.** Isoforms consisting of either exon 1a, b, b’ or the novel exon 1c followed by only canonical main exons. **B.** Isoforms starting with exon 1a or b followed by canonical exon 2 to exon 12 and then followed by a novel exon 13b and novel exon 14 (new C-Terminal end). **C.** Isoforms with starting exons 1a or b and novel exon 6b, ending in the canonical C-Terminal End (two isoforms) or with a novel exon 13c (one isoform). **D.** N-Terminal Isoforms (known), with either starting with exon 1a, b or b’ and ending preliminary due to an alternative exon 7b. **E.** Novel N-Terminal isoform, shorter than the known (see D; therefore prefixed with ‘s”), ending in a novel exon 3b / exon 4b. This isoform is the shortest isoform in the overall pattern. **F.** Novel isoform group consisting of different splicing events upstream exon 3, resulting in a shift of the start codon inside exon 3 with valid open-reading frame. Three variants exist, differing in the 5’-end (either canonical C-, novel C- or N-Terminal end).

### Annotation and prediction of functional Domains in known and novel transcripts

After identification of 12 novel and 6 known PGC-1α isoforms we classified and annotated those also with known functional protein domains. We assigned known functional protein domains to the known isoforms and extrapolated those also on the potential novel transcript variants using motif annotation and prediction tools (detailed search strategy and references see supplemental methods) revealing existence of six main protein domains: Two transcription activation domains (AD1, residues 30-40 and AD2, residues 82-95), a protein-recognition motif involved in transcriptional regulatory processes (*LLXXLL*-motif, residues 141–147), the PDB domain 3D24|D involved in binding of the estrogen-related receptor-alpha (ERRalpha, residues 198-218), a binding domain for interaction with the interaction partner PPARγ (residues 292–338), and finally a RNA-Binding-motif, possibly involved in splicing processes of downstream mRNA targets (residues 677–746).

Integrating the structural differences on mRNA sequence level as well as annotation of functional domains led to separation of 6 groups of isoforms:

The first group characterized by a differing starting exon in each isoform, is built of 13 exons, encoding all known protein domains, and leading to protein lengths between 793 and 802 amino acids (aa) (Figure 3, A). This group includes the “reference” long canonical isoform PGC-1α-a as well as two more known (PGC-1α-b and PGC-1α-c) and a novel isoform (named PGC1 α-d). The latter contains a novel starting exon which we denominate exon 1c, according to the known starting exons 1a, 1b and 1b’).

The second group of two novel isoforms includes transcripts encoding all known PGC-1α protein domains (810 and 814 aa) and can be classified by a novel C-terminal end, built by alternative splicing events in exon 13 and the former 3’-UTR (Figure 3, B). They consist of either exon 1a or 1b followed by canonical exons 2-12 and end with two novel exons (exon 13b in CT-PGC-1α-a and exon 14 in CT-PGC-1α-b).

The third group consisting of three novel isoforms, which contain an alternative (shorter) exon 6b with preserved open-reading frame (Figure 3, C). In this group, no known protein domains are affected. Two isoforms end at the canonical C-terminus (PGC-1α-trE6-a and PGC-1α-trE6-b, 785 and 781 aa), includes a novel exon 13c, resulting from a splice event between exon 13 and the 3’-UTR (PGC-1α-trE6-E13c, 783 aa).

The fourth group consists of the known N-terminal Isoforms with premature stop within a truncated exon 7 (NT-PGC-1α-a, NT-PGC-1α-b and NT-PGC-1α-c; 257-270aa), missing the PPARγ-binding domain as well as the RNA recognition motif (Figure 3, D).

A fifth group with a short single isoform (sNT-PGC-1α, 117 aa), built up by exon 1a, exon 2 and two novel exons (3b and 4b), only contains the two activation domains AD1 and AD2 but is missing all other functional domains (Figure 3, E).

Finally, the last group of (novel) isoforms is characterized by splicing events upstream of exon 3, resulting in a shift of the start codon inside exon 3 with valid open-reading frame (Figure 3, F). Here, three variants exist (PGC-1α-E3c, CT-PGC-1α-E3c and NT-PGC-1α-E3c), differing in the 5’-end (either canonical C-, novel C- or N-terminal end). While all of them lost the activation domains AD1 and AD2, only the NT-PGC-1α-E3c misses also the PPARγ-binding domain as well as the RNA recognition motif.

### Discovery of a novel promoter region of PGC-1α with high conservation in murine and human heart

The novel transcript PGC1a-d in the murine dataset contains a new starting exon (exon 1c; Figure 3A), yet otherwise exhibits the same sequence as the known PGC-1α-a (Figure 3, A), which is the major and most abundant isoform in our and in all published datasets. Exon 1c originates downstream of the known promoter region for exon 1a (Figure 4, A). This finding suggests, particularly considering the known mechanisms of transcription initiation within this gene locus, identification of a new promoter site. Using promoter prediction tools we identified a transcription initiator as well as a promoter element preceding the novel exon 1c, both necessary elements for mammalian transcription start sites (Figure 4, B). Remarkably, we found a homologous transcript with the new exon 1c in multiple Full-Length Reads within the human SMRT-Dataset as well. Additionally, there seems to be high conservation across mice and men, as it shares most of the predicted amino acids. For analysis of tissue-specific expression of the Exon1c-transcript, we designed primers covering the exon 1c-exon2 junction and confirmed sequence identity of the amplified PCR fragment by direct sequencing (Supplemental Figure S02, F). This novel Exon1c-transcript is expressed in different murine organs, with highest expression in brown adipose tissue and skeletal muscle (Figure 4, C).

**Figure 4:**
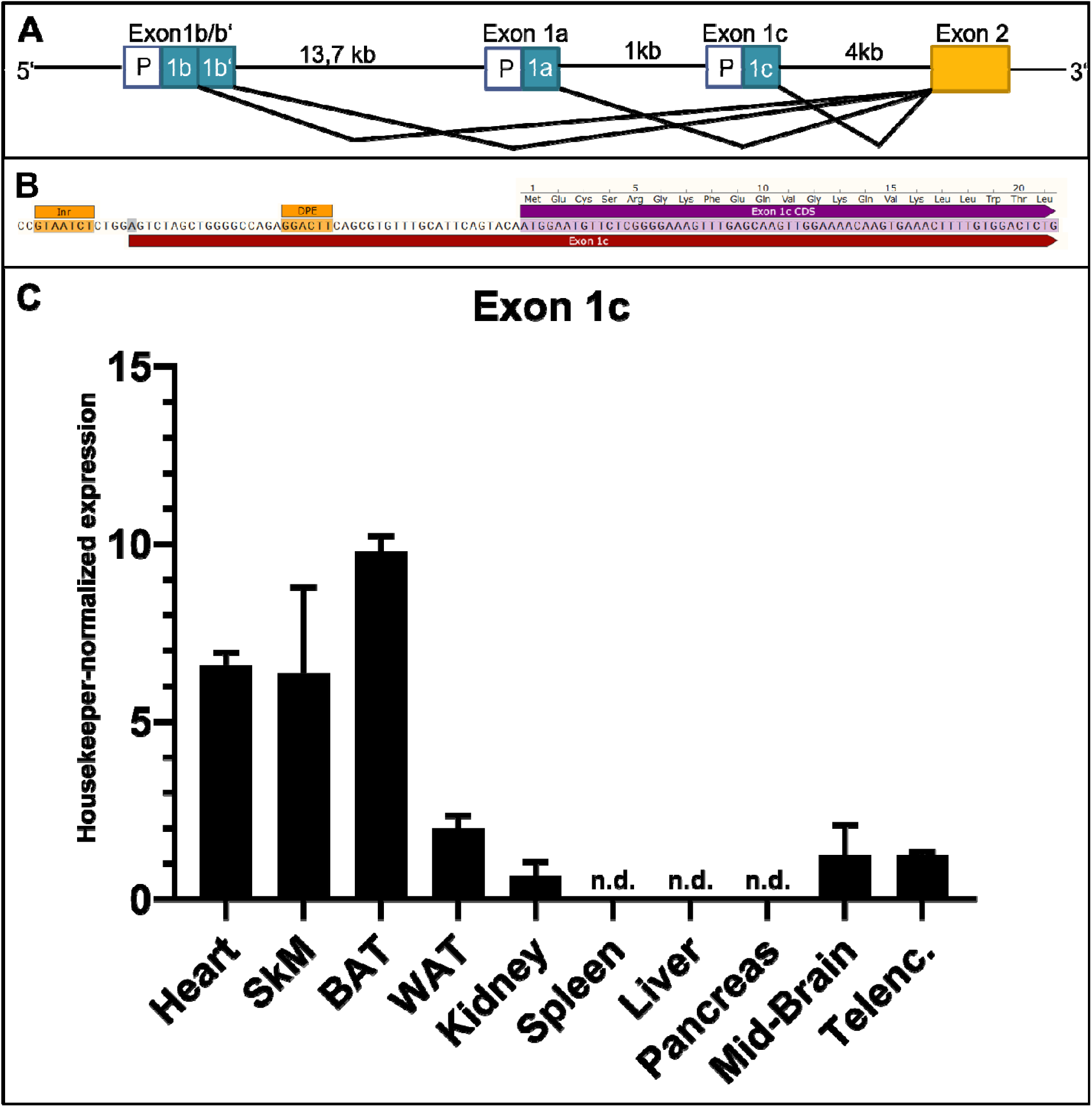
Organ-specific Expression profile of E1c-transcript (Exon 1c) **A.** Graphical illustration of PGC-1α-promoters and corresponding starting exons. The canonical promoter, responsible for exon 1a, lies between the alternative (known) promoter (regulating exon 1b and b’) and the new putative promoter site controlling novel exon 1c. **B.** Novel predicted promoter site associated to exon 1c. Prediction (using ‘ElemeNT’^57^) of mammalian <u>In</u>itiato<u>r</u> element (Inr) and corresponding <u>D</u>ownstream <u>P</u>romoter <u>E</u>lement (DPE) around the Transcription Start Site (marked in grey) of exon 1c**. C.** Expression data of DNA-samples derived from qPCR with primers targeting new Exon1c-Exon2-Junction in heart, skeletal muscle (SkM), brown adipose tissue (BAT), white adipose tissue (WAT), kidney, spleen, liver, pancreas, mid-brain and telencephalon (Telenc.), normalized to housekeeper (NUDC) and factorized by 1000 for better visualization. Highest Expression can be observed in BAT and muscle tissue (heart, SkM); lower, but detectable, expression levels in kidney, WAT and brain. No detection (n.d.) of the new junction could be seen in liver, pancreas and spleen.

### Diet-induced obesity (DIO) leads to lowered alternative promoter driven expression

Gene regulation is accomplished by a variety of mechanisms including differential promoter usage. Thus, we analysed to what extent the different promotors were used for expression of PGC-1α using qPCR-primer-sets covering specifically each starting exons (table S01). Additionally, we were interested in metabolic effects on expression of PGC-1α. Thus, we investigated expression PGC-1α in metabolically dysregulated/challenged mice that were fed a high-fat-high-sucrose diet, leading to diet-induced obesity (DIO) and a prediabetic state (Figure S04).

Notably, for exon 1a (under the control of the canonical promoter) as well as exon 1c (under the control of the novel promoter), comparison of direct expression levels between DIO and lean mice were not showing differences (Figure 5, A). On the other hand, exon 1b and exon 1b’, originating both from the alternative promoter, were significantly lower expressed in DIO compared to lean mice (fold changes: exon 1b 0.71, exon 1b’ 0.36 with corresponding p-values of 0.03 and 0.04, resp.).

**Figure 5:**
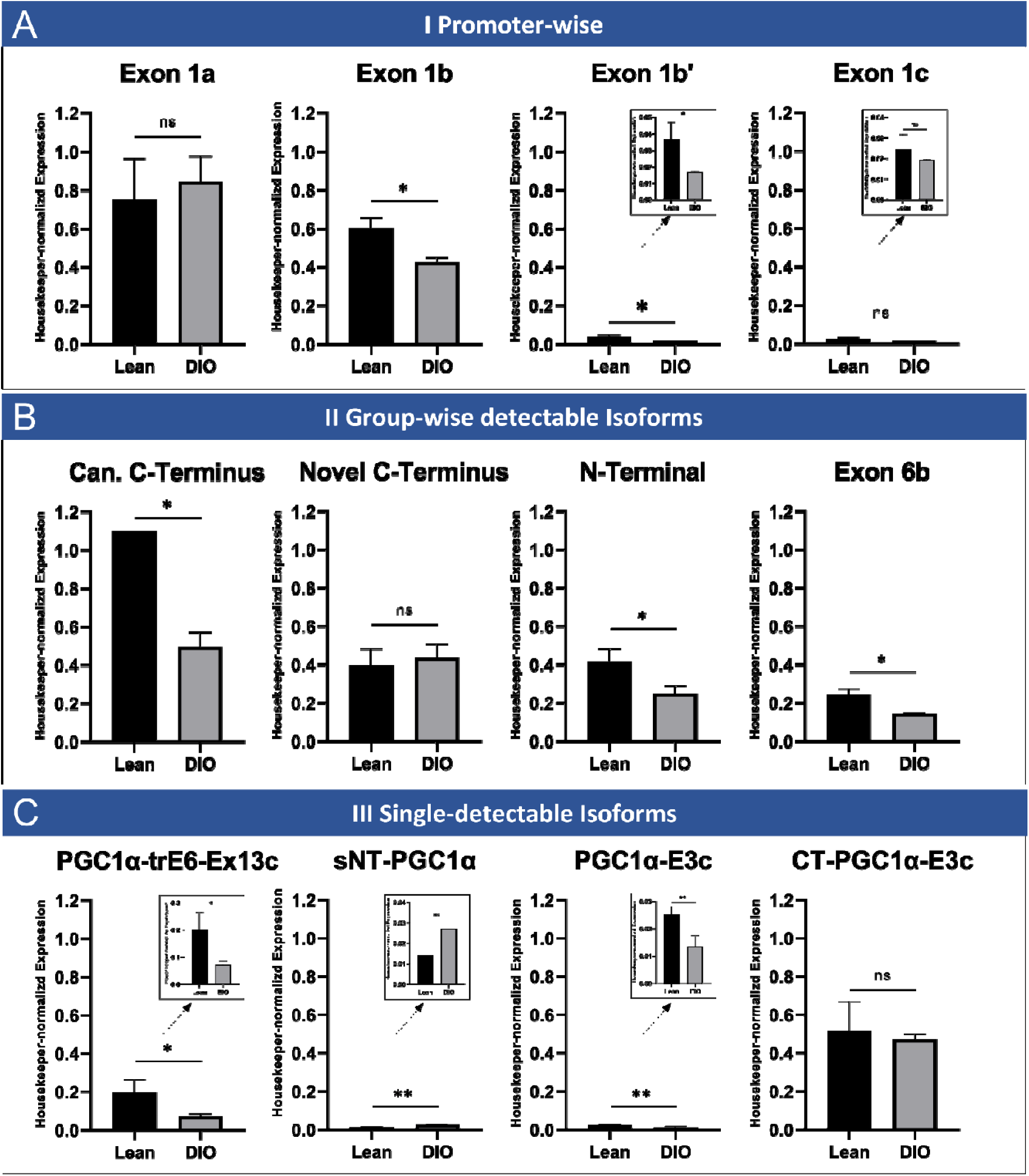
PGC-1α isoform expression in heart in lean or diet-induced obesity (DIO) mice. Housekeeper-normalized expression levels at baseline for either lean mice (black bars) or mice with diet-induced obesity (DIO, grey bars). Shown is isoform expression for PGC-1α starting exons (**A**), for the four group-wise testable isoforms (**B**) and for the four single-detectable isoforms (**C**). **A.** Isoform expression by promotor usage. While expression levels for exon 1a and exon 1c are similar between both conditions, transcripts containing exon 1b and exon 1b’ are significantly lower expressed in DIO. **B.** Expression of group-wise detectable isoforms. While isoforms with novel C-terminal end are unchanged between both diets, the other group-wise detectable isoforms show significantly lower expression in DIO. **C.** Expression for isoforms which are single-detectable through PCR. While CT-PGC-1α-E3c is equally expressed under both conditions, the other single-detectable isoforms show significantly reduced expression in DIO. Data acquired by qPCR using cDNA using primers covering specific starting exon 1 (a, b, b’ or c) and exon 2 resp. using primers covering specific exon-exon-junctions. Expression values are normalized to housekeeper NUDC. n=4 each, bars depict mean values, error bars represent SD. Significance calculated by unpaired student’s t-test (ns p>0,05, *p≤0.05, **p≤0.01, ***p≤0.001).

### PGC-1α expression is reduced in DIO except for novel C-Terminal Isoforms, CT-PGC-1α-E3c and sNT-PGC-1α

Next, we investigated the influence of high-fat diet on expression levels of the different isoforms. Those isoforms which can be detected solely by covering common junctions (canonical-C terminus, novel C-terminus, shorter N-terminus, truncated Exon 6b’), showed significantly lowered expression under DIO with fold changes in between 0.47 and 0.6 (Figure 5, B), p-values between 0.02 and 0.05, resp.), with exception of the novel C-Terminal Isoforms (p-value 0.97). Second, in those isoforms with ability to be detected uniquely by PCR (Figure 5, C), a more heterogenous pattern can be observed: PGC-1α-trE6-Ex13c as well as PGC-1α-E3c are also significantly lower expressed in DIO (fold-changes 0.36 and 0.53, p-values 0.006 and 0.02, resp.). CT-PGC-1α-E3c on the other hand is equally expressed under both conditions, and sNT-PGC-1α shows slightly but significantly higher expression in DIO compared to lean mice (fold-change 1.9, p-value 0.006).

### Expression of PGC-1α is reduced 3 days but partially recovers 16 days after I/R in the infarcted area

After proving evidence for DIO-associated differential PGC-1α isoform expression, we aimed to analyse potential (dys-)functional regulation of the PGC-1α transcriptome in cardiac ischemia/reperfusion (I/R) injury serving as experimental model for myocardial infarction. We used a well-established protocol and compared samples at baseline as well as 3 and 16 days post I/R from the infarcted area and remote area to sham-operated controls (workflow see Figure 1, B).

Under basal conditions in murine heart, all starting exons and isoforms are expressed in similar level (Figure 6 A-D). I/R-injury leads to downregulation of all starting exons 3 days post intervention in the infarcted area compared to sham-control (0.32- to 0.38-fold, *p<0.05 for exon 1a, 1b’ and c and ***p<0.001 for exon 1b). In contrast, in the remote area the expression levels between sham and I/R are equal.

**Figure 6:**
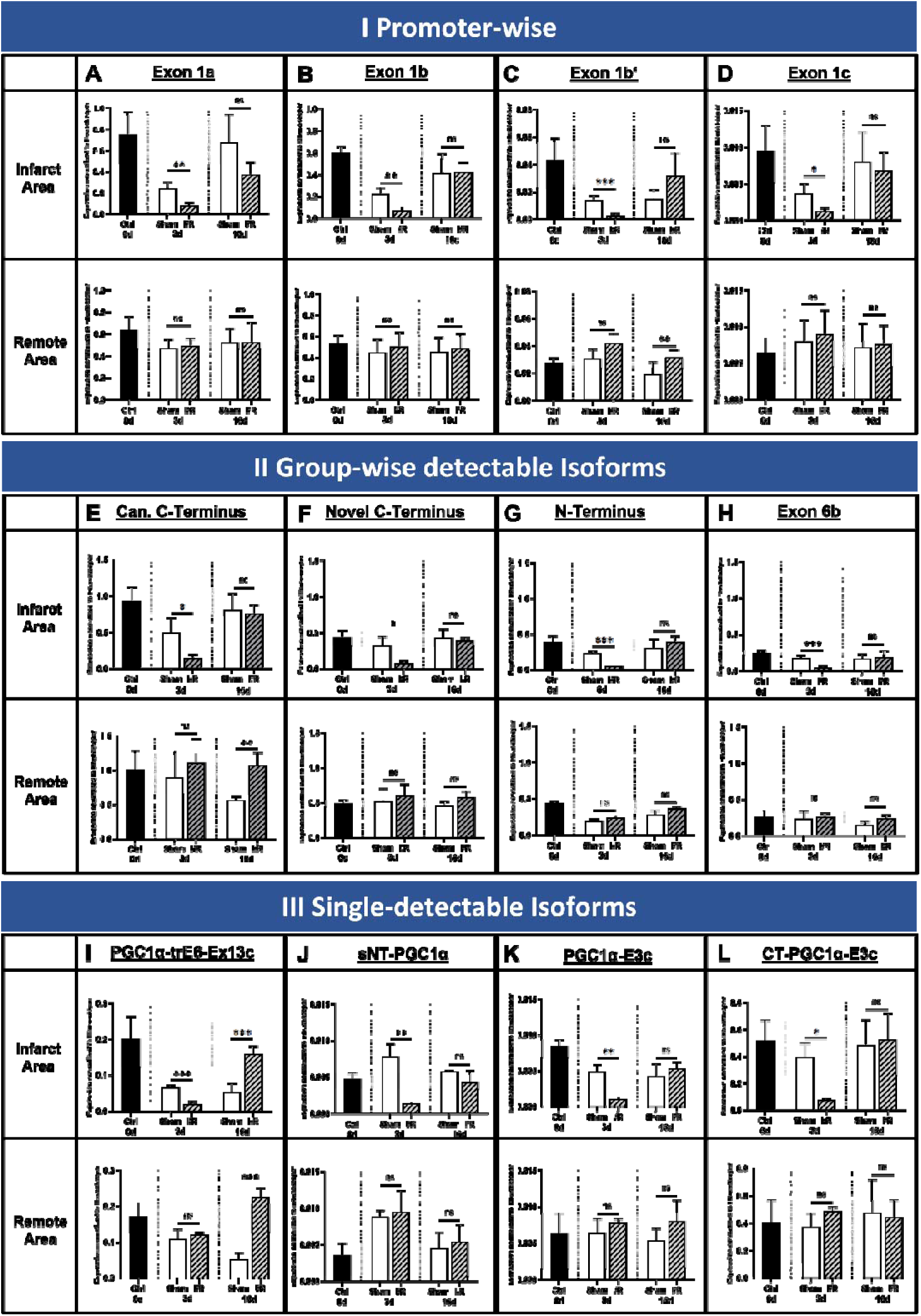
PGC-1α isoform expression in heart at baseline and 3- and 16 days post I/R. Expression levels at baseline (black bars) and 3- or 16-days post sham-surgery (sham, white bars) or Ischemia / Reperfusion injury (I/R, striped bars) for different PGC-1α starting exons (A-D), for the four group-wise testable isoforms (E-H) and for the four single-detectable isoforms (I-L). Housekeeper-normalized expression values in infarct area (upper row) and remote area (bottom row). **A.** Exon 1a, originating from the canonical promoter. **B.** and **C.** Exons 1b and 1b’, under control of the alternative (known) promoter. **D.** Novel Exon 1c, originating from a new promoter site. **E.** Isoforms with canonical C-terminal ending. **F.** Isoforms with novel C-terminal ending. **G.** N-terminal isoforms. **H.** Isoforms with the novel exon 6b. **I.** PGC-1α-trE6-Ex13c **J.** sNT-PGC-1α **K.** PGC-1α-E3c **L.** CT-PGC-1α-E3c. I/R leads to downregulation of canonical and novel PGC-1α starting exons as well as most of the single- and groupwise-detectable PGC-1α-Isoforms 3d post I/R, suggesting a general downregulation of PGC-1α in infarct area 3d post I/R. At 16 days post I/R, a recovery can be observed. Moreover, PGC-1α-trE6-Ex13c is showing ‘overcompensative’ behavior in infarcted and remote area. In all other isoforms, the expression in the remote area is not affected, neither at day 3 nor 16d post infarction. Detailed description see text. Data acquired by qPCR using cDNA using primers covering specific starting exon 1 (a, b, b’ or c) and exon 2 resp. using primers covering specific exon-exon-junctions. Expression values are normalized to housekeeper NUDC. n=4 each, bars depict mean values, error bars represent SD. Significance calculated by unpaired student’s t-test (ns p>0.05, *p≤0.05, **p≤0.01, ***p≤0.001).

During recovery, 16 days post I/R, almost no difference is observed between I/R and sham (Figure 6, A-D). This is due to a significant increase of expression in the infarcted area from 3 to 16 days post I/R (upper rows; fold change exon 1a 4.73, exon 1b 5.85, exon 1b’ 12.88, exon 1c 5.56; corresponding p-values exon 1a 0.013, exon 1b 0.016, exon 1b’ 0.04, exon 1c 0.02). In contrast to that, the expression in the remote area over time stays unchanged (bottom rows).

Additionally, we were also interested if changes in expression also occur the group-wise or single-detectable isoforms. Here, a similar pattern as for the promotor variants was observed also for most of the splice variants: When compared to sham, I/R induces a transient decline of transcripts in the infarcted area on day 3 post I/R, which was followed by recovered expression on day 16. Expression in the remote myocardium was not altered except for the C-terminal Isoforms and PGC-1α-trE6-Ex13c (Figure 6, E-L; fold changes for the infarcted area ranging from 3.08 to 8.13, p-values from 0.0001 to 0.03).

### I/R in DIO leads to heterogenous pattern of PGC-1α-expression

As both DIO and I/R individually changed PGC-1α expression in cardiac tissue, we were curious if combining both hits would have an additional effect. Therefore, I/R was repeated in mice fed 10 weeks with high-fat diet, using the same protocol which was used for each modality separately before (see scheme in Figure 1, B). At baseline, the majority of all PGC-1α transcripts exhibit lower expression levels in DIO than lean mice (Figure 7, A, Figure 5 and color-coded Figure S07). Exceptions are CT-PGC-1α-E3c (Figure 7, horizontally half-filled diamond-shape) and isoforms with the novel C-Terminus (Figure 7, empty square), which are equally expressed, and transcripts with exon 1b’, which are higher expressed in DIO. Three days post I/R (Figure 7, B, and color-coded Figure S07), in the infarcted area this changes as, beside low absolute expression values in both groups, all transcripts are either equally or higher expressed in DIO. This effect appears transient, since 16 days post I/R (Figure 7, C) a similar pattern to baseline can be observed for most transcripts. However, transcripts starting with exon 1a (filled circle) and exon 1b’ (empty circle) show higher expression in DIO than lean mice at this time point, while isoforms with the N-terminal end (vertically half-filled square), are even lower expressed than before (see also Figure S07). Thus, expression distribution at 3 days post I/R seems altered by combination of metabolic and ischemic hit, with incomplete recovery over time in the infarcted area.

**Figure 7:**
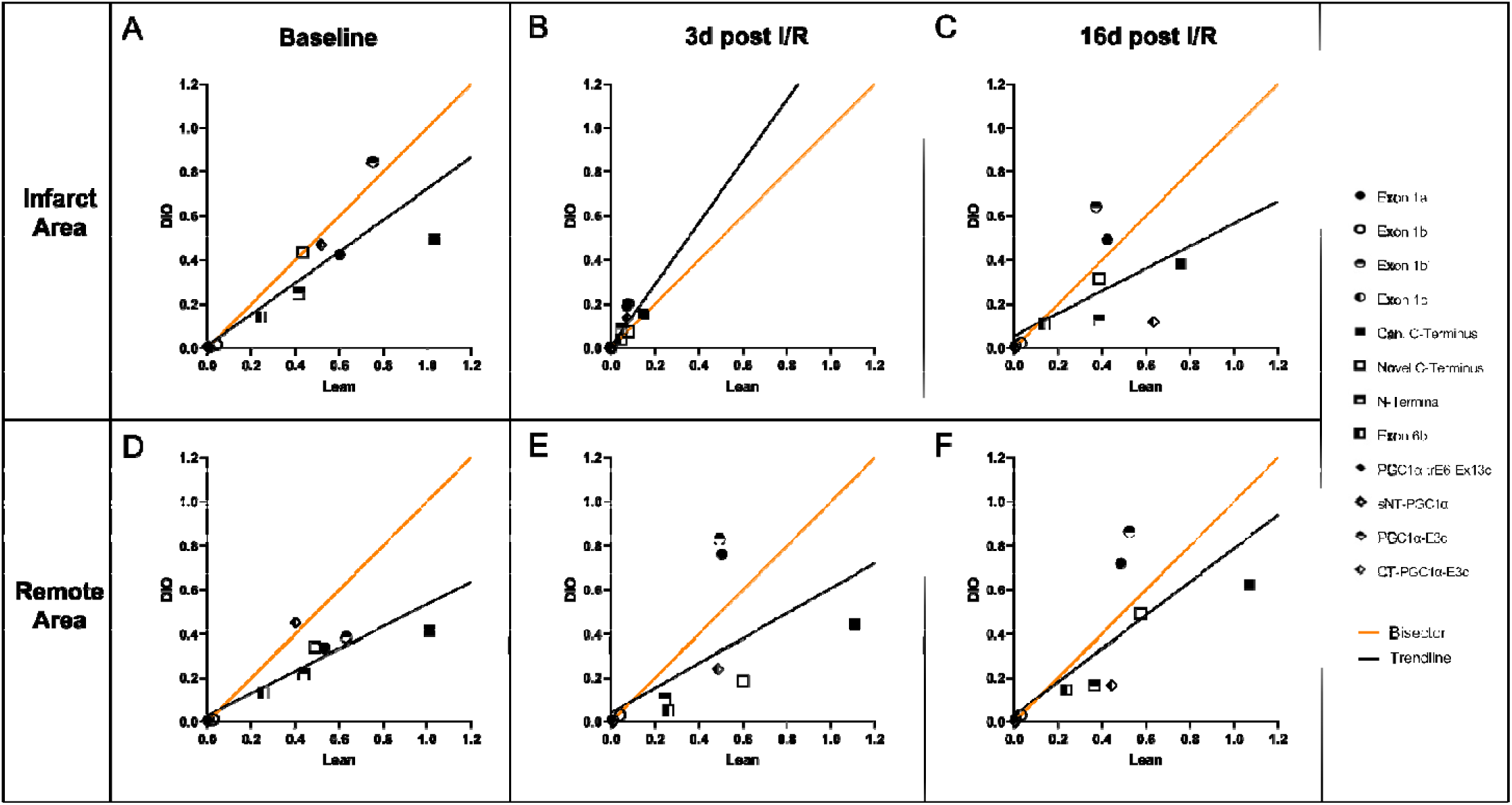
Distribution of PGC-1α isoform expression in lean vs DIO under I/R. Housekeeper-normalized expression levels at baseline and 3- or 16-days post Ischemia / Reperfusion injury as relation between DIO (y-axis) or lean mice (x-axis). Shown are values for the infarcted area (upper row) and remote area (bottom row). Each dot represents average expression value (n=4) of each transcript (either promotor-wise, group-wise or single-detectable isoforms, with same categories as in Figure 5). Black line represents trendline for average expression values of all transcripts. Orange line shows bisector, where values above mean higher and underneath mean lower values in DIO than lean mice. **A.** Baseline expression. The majority of all PGC-1α transcripts exhibit lower expression levels in DIO than lean mice, with few exceptions (see text).**B.** Expression 3 days post I/R. In contrast to baseline, in the infarcted area, all transcripts are either equally or higher expressed in DIO. **C.** Expression 16 days post I/R. In the infarcted area in contrast to remote, the combination of metabolic and ischemic hit leads to incomplete recovery of expression over time.

The remote area shows also lower expression of all isoforms in DIO than lean mice at baseline (Figure 7, D, Figure 5 and Figure S07). Three days as well as 16 days post I/R in the remote area, expression in transcripts with exon 1a (filled circle) and exon 1b’ (horizontally half-filled circle) are significantly higher in DIO than lean mice (Figure 7, E and F; Figure S07), while all other transcripts resemble baseline expression values.

## Discussion

PGC-1α, the most important member of the PGC-1 gene family of transcription factors, is regulated on transcriptional and post transcriptional level. In skeletal muscle, alternative splicing and different promoters generate various PGC-1α isoforms with distinct functions. Mice with cardiomyocyte-specific deficiency of total PGC-1α develop dilatative cardiomyopathy(24), vice versa overexpression of PGC-1α1 enhances contractility(25). Though playing a major role in the heart orchestrating mitochondrial biogenesis(4) and function, little in-depth information about the cardiac transcriptome or possible differential expression PGC-1α is available.

This study aimed to fill this gap by combining latest-generation sequencing with metabolic and ischemic challenges to investigate PGC-1α isoform expression in cardiac tissue.

Here, we report three major findings:

1. For the first time, we generated a reliable, full-length dataset on cardiac transcript expression.
2. We analysed expression pattern of PGC-1α isoforms and identify novel and known PGC-1α transcripts being differentially expressed under high-fat diet, a model for diabetic cardiomyopathy.
3. In murine I/R injury, we demonstrate that promoter-driven PGC-1α transcripts are downregulated and partially recover after 16 days post I/R in the infarcted area.

### Deciphering full-length, high-fidelity transcriptome with discovery of 58000 unique isoforms in murine and human heart

In order to address the complexity of splice variants with high-quality long-read sequencing, we performed full-length, high-fidelity transcriptomics of both murine and human heart for the first time. More than 58000 unique, full-length sequences with very good quality were achieved. To our knowledge, full-length transcriptomic data are not yet available for most organisms, especially not for mammals. One earlier study reported even lower number of genes and transcripts in human and mouse brain, but overall comparable quality (30). Using the SMRT technique we add important information to the analysis of the cardiac transcriptome as this method avoids assembly of transcripts from classical short read sequencing that can lead to wrong assumptions on sequence identity.

Naturally, there are always technical limitations regarding sequencing methods in general and the full-length transcriptome in special: First, good pre-analytics is crucial as the quality of the original RNA is limiting the data availability on transcriptomic level. Therefore, only samples with high RNA integrity score (RIN > 9) were used for further analysis. Second, to achieve 99% sequence identity to the true full-length mRNA sequence, automated data analysis removes sequences for which quality control criteria cannot be reached. Therefore, some potentially true sequences get lost throughout the process. To overcome this, we performed, a second, manual readthrough, using the non-filtered, full-length, non-concatemers, after the automatic pipeline assessment, thus minimizing loss of true mRNA sequences. Third, although we do not need to map short reads for assembly, mapping of the long read sequences to the genome for correct assignment of the transcriptomic reads to genomic loci must be performed. The fidelity of the mapping relies on the underlying genomic data (which is in mice and human overall good) and additionally on the algorithm used. To overcome potential errors regarding software issues we used different available mapping tools (minimap2(31), STAR(32) and GMAP(33)) as well as different genomic database sources (GENCODE, ENSEMBL) resulting in comparable results. Fourth, if focussing on PGC-1α, limitations are also that low abundant transcripts are more difficult to detect. Here, long-read sequencing using SMRT can detect one unique transcript even if there is only one single molecule inside the sample. Even using enough so called Zero-Mode Waveguides (each containing the ‘hole’ in which one molecule is sequenced), sensitivity is always limited, and we cannot exclude that some very low abundant transcripts / isoforms of PGC-1α may not be (physically) detectable.

### Investigation of PGC-1α expression pattern leads to discovery of 12 novel PGC-1α isoforms and a novel promoter

Focussing on PGC-1α, our bioinformatic pipeline identified most of the known PGC-1α transcripts, and strikingly, 12 novel isoforms. Furthermore, we discovered a putative novel promoter, controlling expression of a new starting exon, which we called ‘exon 1c’, following the terminology of known PGC-1α promoters. Notably, we identified this promoter also for human heart tissue enabled by existence of conserved transcripts. The novel predicted promoter site associated to exon 1c has a mammalian initiator element (Inr) and corresponding downstream promoter element (DPE) around the transcription start site of exon 1c, raising confidence about the existence of the promotor element.

The newly detected isoforms are in line with regulated gene expression which is commonly considered a combination of alternative splicing events and usage of different promoters: PGC-1α1, PGC-1α-b and PGC-1α-c, all known from literature, share the same ‘sequence body’ but start with either exon 1a, exon 1b or exon 1b‘. In this present study we demonstrated a new transcription start with the novel exon 1c and the very same sequence afterwards (hereafter called PGC-1α-d).

In the same manner, also the novel isoforms are derivatives of known sequences, but result from alternative internal splicing, giving rise to new exonic features within the mRNA sequence. Interestingly, most of those isoforms also follow the principle of shared core sequence with different starting exons. Although we did not perform SMRT sequencing in other organs, we confirmed expression of exon 1c in various organs including heart and skeletal muscle as well as brown adipose tissue by quantitative PCR. Therefore, we suggest that a novel promoter specifically active in those tissues drives expression of exon 1c.

Of special interest is also one group of three novel isoforms lacking ‘classical starting exons’ due to alternative splicing events shifting the open reading frame. All of them share the same novel starting exon 3c, which originates from within the sequence of canonical exon 3. This is of special interest as it resembles the structure of an isoform previously known in human liver (L-PGC-1α(34)), which was not detected in other vertebrates so far. As the sequence is not identical to the human liver isoform, but both share the very same splicing mechanism and structural elements, they seem to be very closely related and are likely functionally similar.

### Pre-diabetic state in mice leads to reduced alternative promoter-driven expression as well as reduction of PGC-1α expression in general

Others and we found that obesogenic diet in mice leads to a prediabetic metabolic state with higher blood glucose and insulin levels, evolving in insulin resistance due to diet-induced obesity (DIO). PGC-1α is functionally implicated in obesity-associated syndromes as high-fat diet counteracts expression of the PGC1 gene family in drosophila(35) and leads to upregulated apoptosis pathways in murine hepatocytes through inhibition of PGC-1α mediated suppression of NFκB(28). We found that DIO is associated with a reduction for most PGC-1α transcripts.

### Transient promoter-driven PGC-1α downregulation in I/R in infarcted area in mice

In the heart total knockout of PGC-1α leads to a DCM-like heart failure phenotype(24). Data on isoform specific functions in the heart are only available for PGC-1α1, suggesting a cardioprotective function when overexpressed(25).

We found that I/R led to a general downregulation of the whole PGC-1α isoform pattern in the infarct zone when compared to sham control, while no effect on the remote area was observed 3 days post intervention. Usually, scar-associated gene programs (like extracellular matrix proteins and inflammatory pathways) are rapidly activated after myocardial infarction(36) in the infarct zone. In contrast to this, PGC-1α-expression in general was downregulated in the infarcted area in our experiments, supporting the theory of a stunned myocardium with reprogrammed metabolic pathways due to lowered energy supply (‘hibernating myocardium”)(37). Thus, transcriptomic regulation of PGC-1α mainly occurs in the infarcted area 3 days post I/R, while expression equalizes to sham controls for all isoforms, indicating a recovery of expression in the subacute phase after I/R injury.

### Combined metabolic and ischemic stress leads to altered PGC-1α expression response

In human cardiac tissue, lipid accumulation and insulin resistance increase vulnerability to ischemia-induced cardiac dysfunction(38). We found a differential expression of PGC-1α isoforms in the remote area uniquely in DIO, which could be explained by a shifted metabolic state in the remote area, while in the infarct area the hibernating myocardium is not capable to do so. Strikingly, when comparing relative expression values directly to those under standard chow, in DIO both the infarcted area and the remote myocardium exhibit downregulated expression of PGC-1α, which is different to lean mice, where only expression in the infarct zone itself is altered. Additionally, one novel isoform, PGC-1α-trE6-Ex13c, depicted a 13-fold (overcompensatory) increase in the infarct zone but no recovery at all in the remote area.

Our working hypothesis is that metabolic changes in diet-induced obesity shift isoforms pattern towards distinct roles, e.g. towards hypertrophy or changes in metabolic pathways. To what extent the newly identified isoforms are cardioprotective or worsen outcome after myocardial infarction, especially under high-fat-diet conditions and prediabetic state, remains to be elucidated. Another open question is hereby the underlying cell type which causes the observed expression changes (e.g. cardiomyocytes, infiltrating immune cells, fibroblasts). Additionally, as direct protein proof of novel isoforms is challenging, we cannot exclude that some of the isoforms are not translated. However, as we selected only sequences with poly-A-tail bioinformatically, and we have confidence on the existence of a transcript because of a full-length sequence, it is very likely that the respective protein is synthesized in vivo.

From a translational perspective, the differences of PGC-1α expression between lean and DIO mice in I/R could be extrapolated on diabetic patients and their unfavorable outcome after myocardial infarction. Future prospective studies, including transcriptomics and metabolomics, are needed to elucidate pathomechanistical insights into the complex regulatory mechanisms PGC-1α expression and function in the heart of those patients.

## Methods

### Mice

Male 12-week-old C57BL/6J mice (Janvier Labs, Le Genest-Saint-Ile, France) were used for all mice experiments. Mice used for functional testing of isoforms underwent myocardial ischemia/reperfusion injury (I/R) or sham-surgery as described previously(39) (see supplemental methods for details). For experiments involving metabolic changes by diet-induced obesity, mice were fed 10 weeks before I/R with a high-fat-high-sucrose diet (24% Sucrose, 60 kJ & Fat; ID S7200-E010; ssniff Spezialdiäten GmbH, Soest, Germany). All other mice were fed with normal chow.

Permission for animal experiments was granted by the Landesamt für Natur, Umwelt und Verbraucherschutz (LANUV) Nordrhein-Westfalen, Aktenzeichen 81-02.04.2017.A401.

### Human tissue

Fresh tissue from human left ventricle, gained as remnant of left-ventricular assist-device (LVAD)-implantation, was aseptically obtained at the University of Duesseldorf cardiovascular department (Institutional Review Board approval 5263R/2015104434), following the Declaration of Helsinki Principles. Directly after surgery, the heart sample underwent deep freezing in liquid nitrogen and was then immediately used for RNA-isolation as described below.

### RNA isolation

Total RNA was isolated using the Fibrous Tissue RNeasy Mini Kit (Qiagen) according to the manufacturers protocol. Quality control and concentration measurements were performed using Nanodrop and 2100 Bioanalyzer (Agilent). Only Samples with RNA Integrity Number (RIN) above 9 were used.

### Library preparation

Total RNA from murine wildtype total heart and human heart sample from left ventricle was isolated as described above and used for library preparation according to the manufacturers protocols (‘Iso-Seq™ Template Preparation for Sequel® Systems) without size-selection.

### Long-read RNA-Sequencing

Long-read Sequencing was performed at the Genomics & Transcriptomics Laboratory of the Biological-Medical Research Center (BMFZ, Heinrich Heine University Düsseldorf) using the Sequel I System by Pacific Biosciences (SMRT), according to the manufacturers protocols. For each murine and human Sample, 4 Single-molecule-Real-Time (SMRT)-cells were used (=4x10^6^ zero-mode waveguide holes per sample).

### Bioinformatic Analysis of SMRT-Data

Bioinformatic analysis used a modified IsoSeq3-pipeline followed by an own pipeline of open-source-tools and scripts (see also supplements). Identification of novel and canonical PGC-1α isoforms used the automatic pipeline followed by similarity search vs. exonic sequences within the refined full-length (FLNC) reads and clustered high-quality-transcripts. Obtained isoforms were then manually curated and annotated according to the canonical nomenclature of known PGC-1α isoforms, followed by in silico prediction of open-reading frames. Running environment of all Scripts was CentOS 7.7.1908-based at the High-Performance Cluster (HILBERT, Centre for Information and Media Technology (CIM/ZIM); Heinrich Heine University, Düsseldorf). Annotation of functional domains in transcripts was performed using open-accessible online prediction tools (see also supplements).

### Primer Design and Sanger-Sequencing for Validation of PGC-1α isoform sequences

Whereas possible, primer pairs (supplemental Tables S01 and S02) were designed to target specifically novel, unique exon-exon-junctions or UTR-exon-junctions within the identified PGC-1α transcripts from long-read sequencing data, followed by quantitative Real-Time-PCR (qPCR) and classical Sanger-Sequencing for confirmation and validation. In addition, qPCR-primers to detect the known isoforms and starting exons were designed (Figure 1 and tables S01 and S02).

### cDNA-Synthesis and quantitative RT-PCR

Total RNA was isolated from heart tissue as described above. qPCR was performed on the Step-One Plus real-time PCR system (Applied Biosystems) with Maxima SYBR Green and ROX qPCR Master Mix (Thermo Scientific). Transcript quantities were normalized to Nuclear Distribution Protein C (NUDC) mRNA. Bioinformatic analysis used a modificated X0-Method approach, which is based on the comparative threshold cycle (CT) method(40).

### Data Availability Statement

The transcriptomic data underlying this article are subject to an embargo of 12 months from the publication date of the article. Once the embargo expires, the data will be available through GenBank (NCBI) / Sequence Read Archive (SRA)] at https://www.ncbi.nlm.nih.gov/genbank/, and can be accessed with accession ID SRR16352840 (mus musculus) and SRR16352587 (homo sapiens).

## Acknowledgments

Computational support and infrastructure were provided by the ‘Centre for Information and Media Technology’ (ZIM) at the University of Duesseldorf (Germany). This work was supported by the Genomics & Transcriptomics Laboratory of the Biological-Medical Research Center (BMFZ) of the Heinrich Heine University Düsseldorf.

## Supplemental Figures & Tables

**Figure S01:**
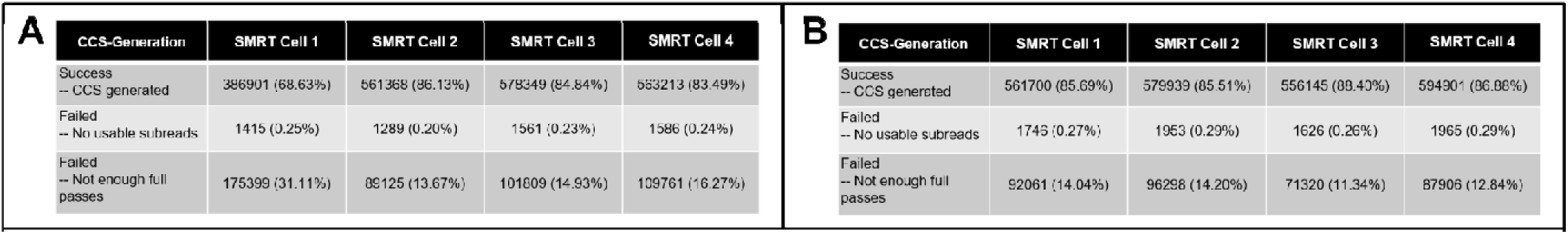
Quality Control for Primary, Secondary and Tertiary Analysis of SMRT-Sequencing. Quality control outputs from different steps of analysis of human (A and C) and murine (B and D) datasets. **A. and B.** Isoseq3-Pipeline Step 1 (Circular Consensus Reads).

**Figure S02:**
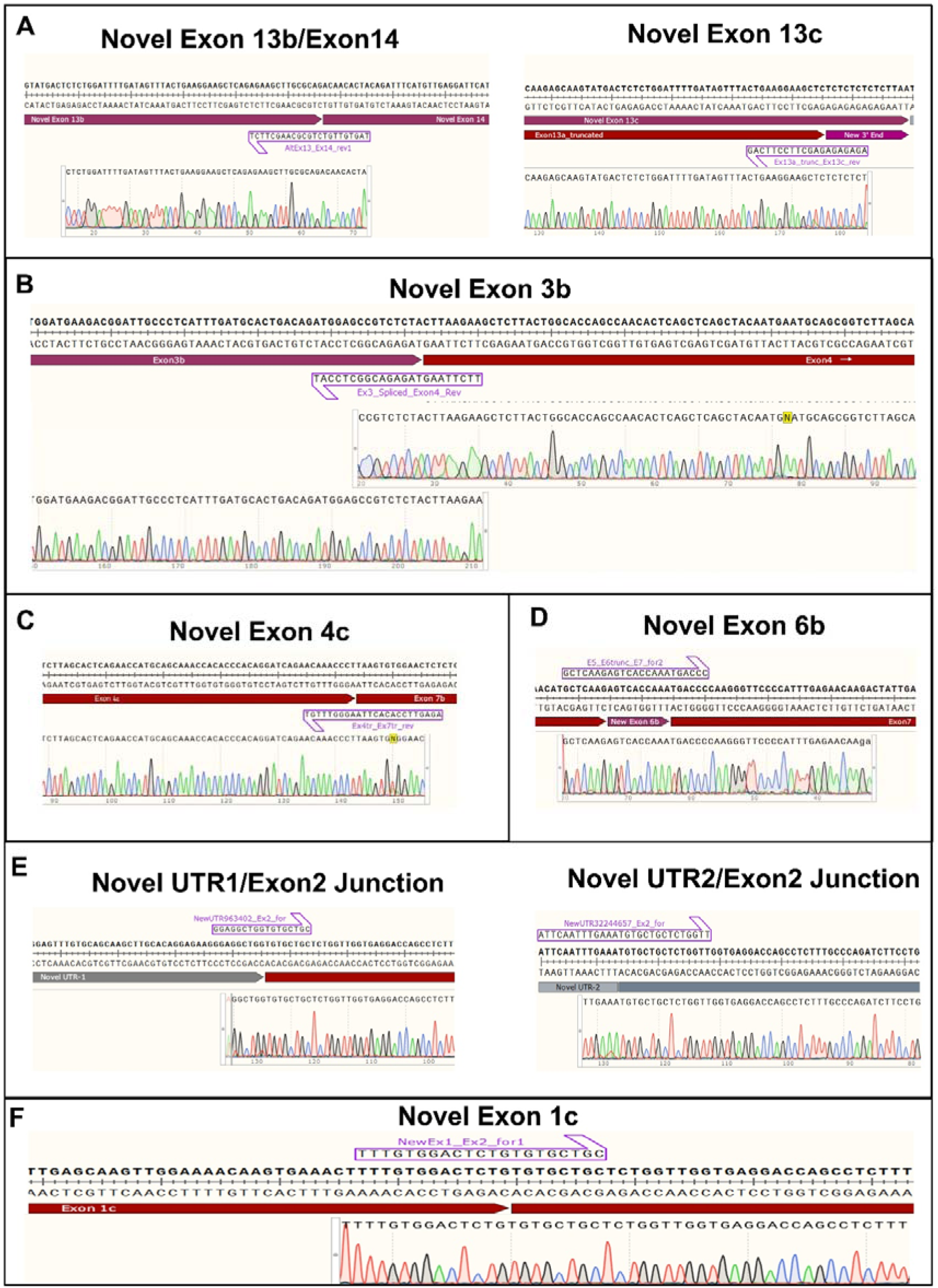
Sanger-Sequencing Results. Shown is the exemplary results of sanger sequencing of fragments after PCR with specific primers (see tables S01 and S02) followed by gel ectrophoresis and extraction. **A.** Novel Exon 13b / Exon 14 junction (left side) as well as Novel Exon 13c (right side). **B.** Junction covering novel Exon 3b/Exon4-junction. **C.** novel Exon 4b / Exon 7b junction. **D.** Junction covering Exon5/Novel Exon 6b/Exon7-Junction. **E.** Novel UTR1-Exon2-Junction, which is unique for PGC1α-E3c (left side) and novel UTR2-Exon2-Junction, which is unique for CT-PGC1α-E3c (right side). **F.** Junction covering the novel Exon 1c and Exon 2.

**Figure S03:**
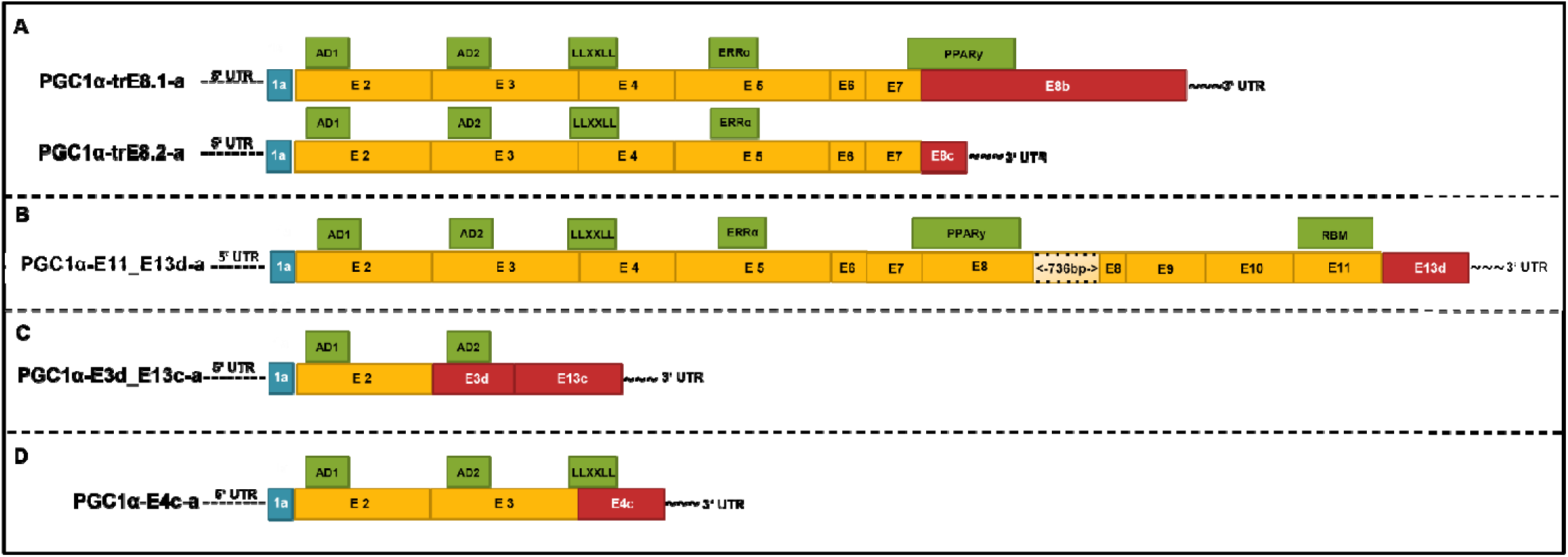
PGC1α-Isoforms in murine heart (failed QC) Graphical illustration of novel, potential PGC1α Isoforms (mRNA) after SMRT-Sequencing which failed QC-Filtering (see Figure 1): Starting exon 1a (blue boxes), canonical main exons (orange boxes), novel/altered exons (red boxes) and functional domains (green boxes, details see text). Length of boxes indicates relative length of nucleotides (true to scale, with exception of exon 8: shortened bp marked). **A.** Isoforms consisting of two versions of a truncated exon 8 with new stop codon within. **B.** Long isoform with skipped exon 12 and novel exon 13d. **C.** Short isoform with novel exon 3d followed by novel exon 13c. **D.** Short isoform with novel exon 4c with stop codon directly afterwards (part of former exon 7)

**Figure S04:**
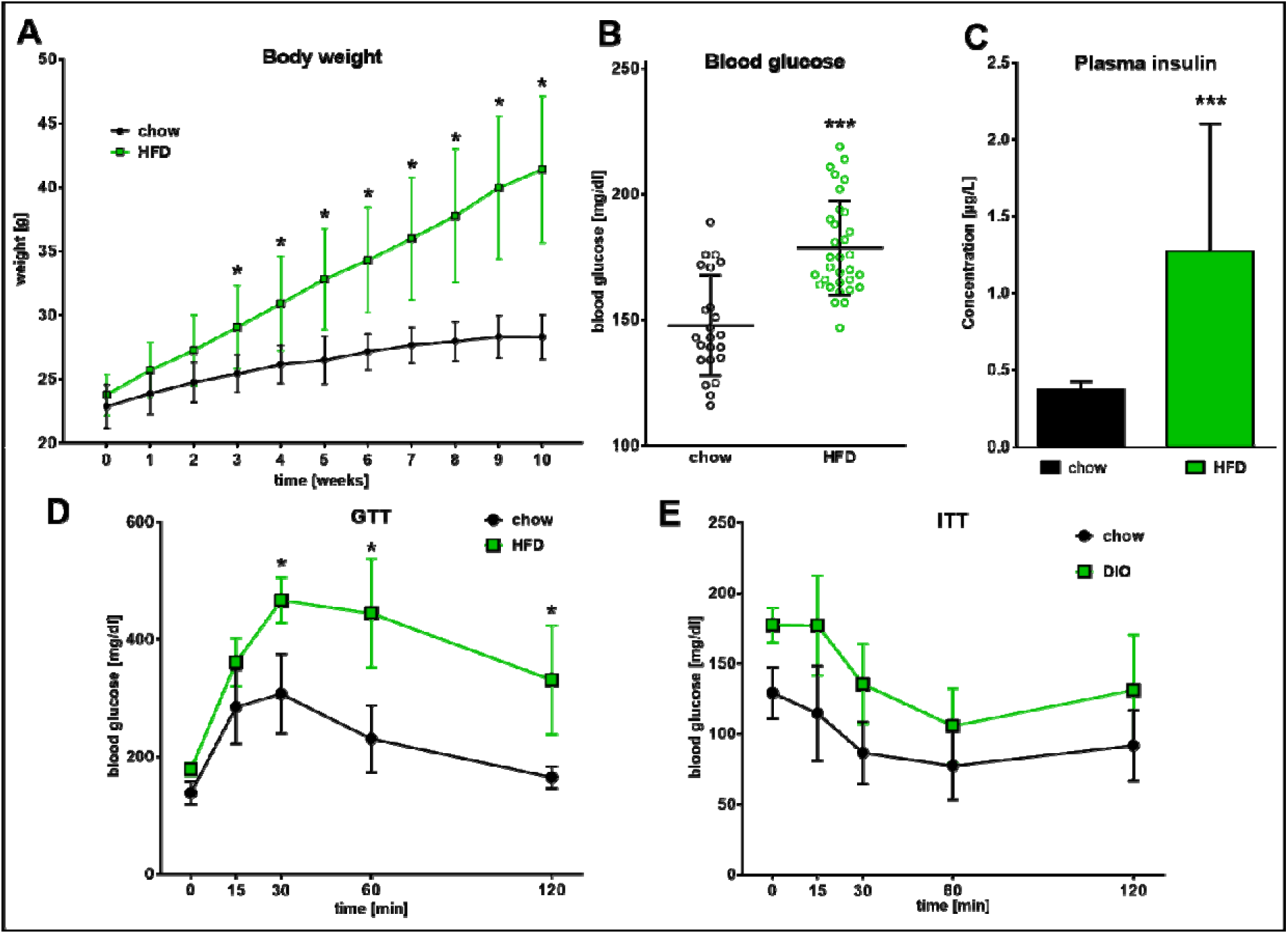
Pre-diabetic phenotype. **A.** Summarized data for body weight development within 10 weeks of standard chow diet (SD) and High-Fat Diet (HFD), n=11. **B.** Summarized data for blood glucose of Ctrl & DIO animals after 9 weeks of feeding (n=23/32). **C.** Summarized data for plasma insulin ELISA of Ctrl & DIO animals after 10 weeks of feeding (n=15/16). **D.** Summarized data for glucose tolerance test of Ctrl & DIO animals after injecting 2 mg glucose per gram body weight after 10 weeks of feeding (n=7/8). **E.** Summarized data for insulin tolerance test of Ctrl & DIO animals after injecting 0,75 U insulin per kg body weight after 10 weeks of feeding (n=11/12).

**Figure S05:**
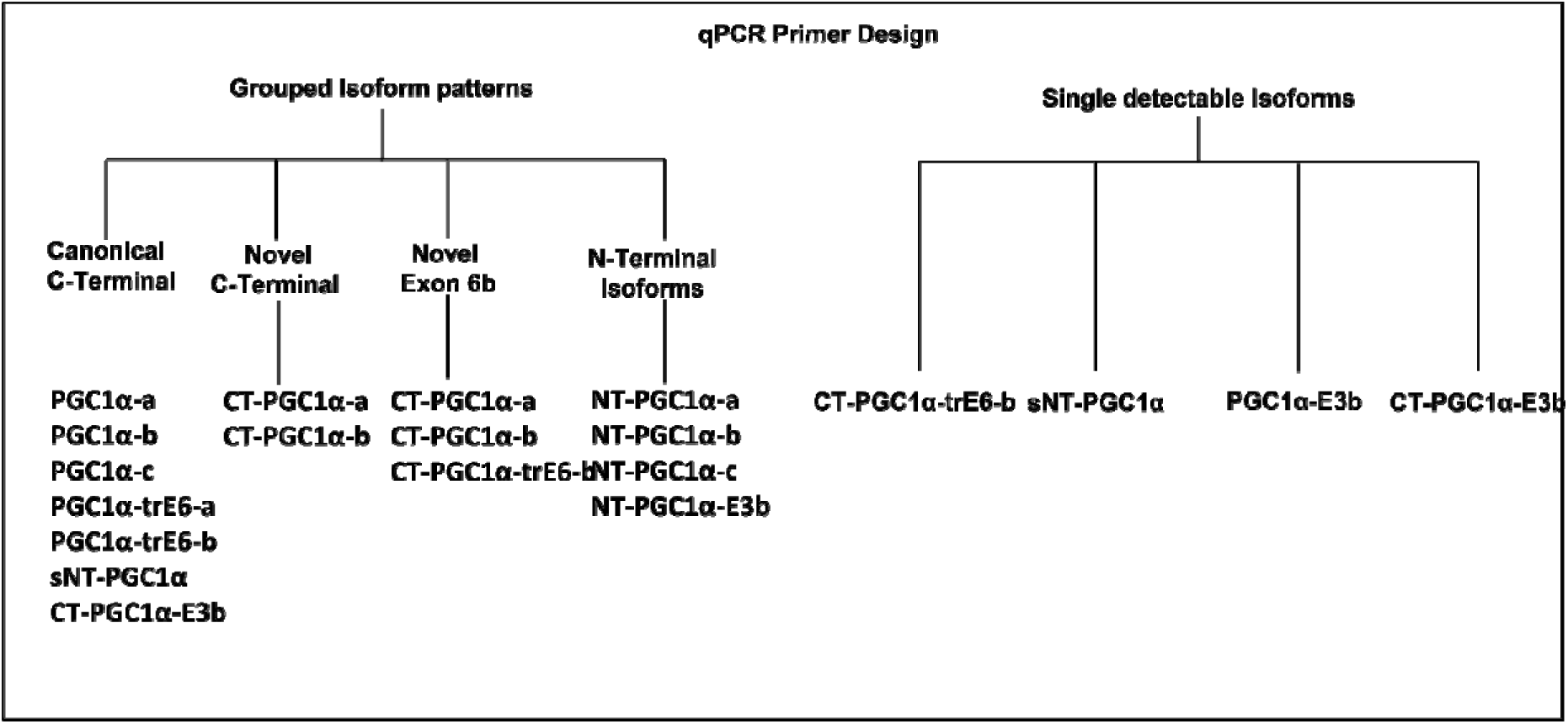
Strategy for Detection of PGC1α-Isoforms in qPCR.

**Figure S06:**
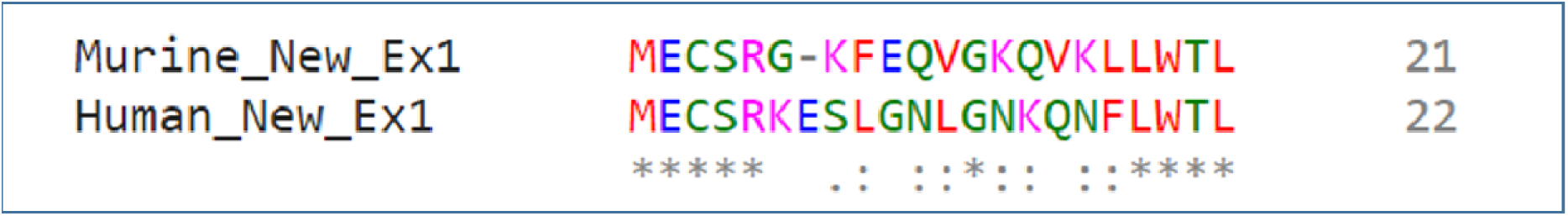
Predicted Open reading Frame Ex1c (murine and human)

**Figure S07:**
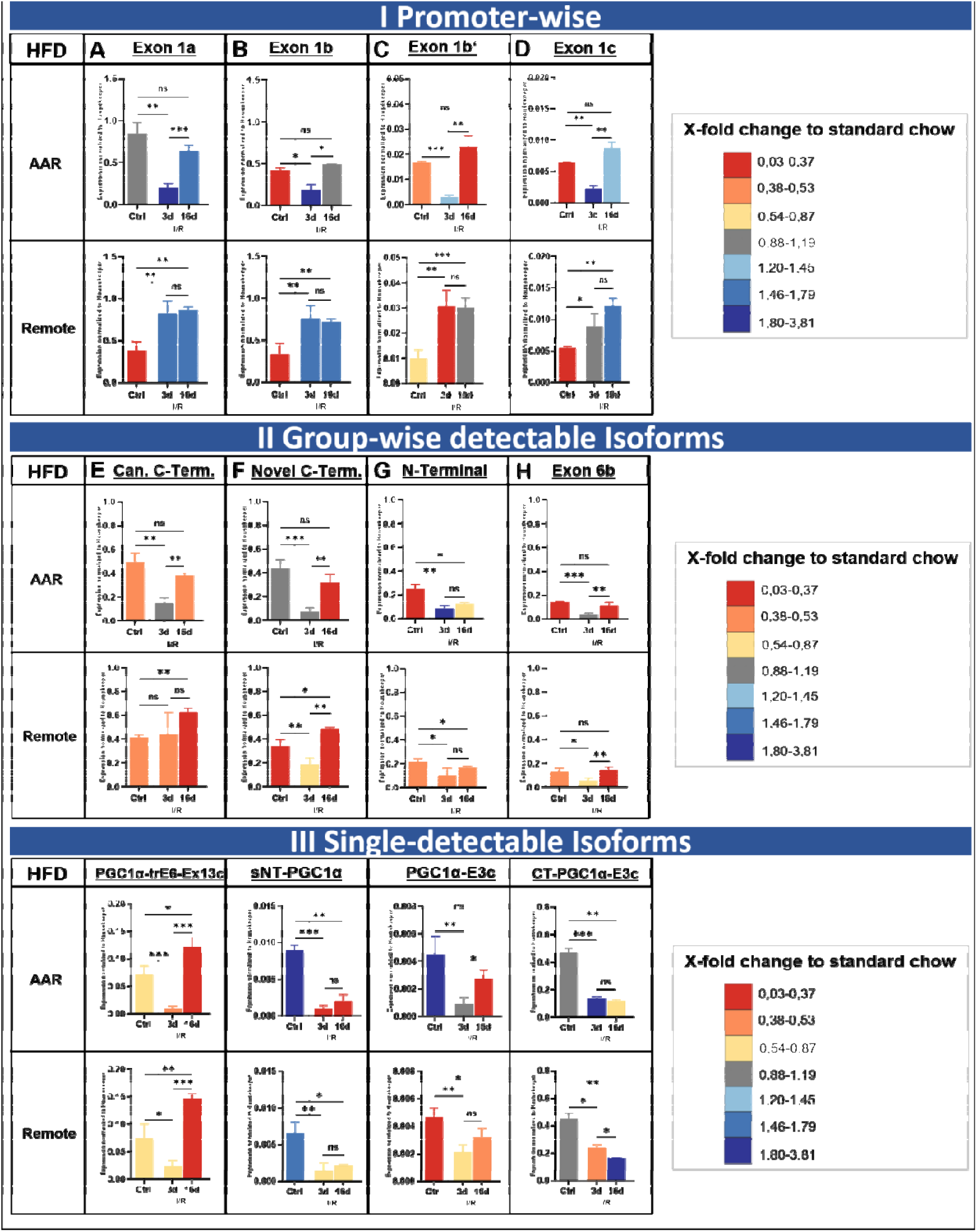
Impact of High-Fat-Diet on Promoter-specific Expression. Expression levels for diet-induced obesity at baseline (Ctrl) and 3- or 16-days post Ischemia / Reperfusion injury (I/R) for different PGC-1α starting exons (A-D), for the four group-wise testable isoforms (E-H) and for the four singe-detectable isoforms (I-L). Housekeeper-Normalized Expression Values in Area-at-Risk (upper row) and Remote Area (bottom row). Colors of bars represent fold-change difference between high-fat (HFD) compared to standard chow (SD) diet, color scheme likewise to heat map illustrations (i.e. dark green means higher, dark red means lesser expression in HFD compared to SD). **A.** Exon 1a, originating from the canonical promoter. **B.** and **C.** Exons 1b and Exon 1b’, under control by the alternative (known) promoter. **D.** Novel Exon 1c, originating from a new promoter site. **E**. Isoforms with canonical C-Terminal ending. **F**. Isoforms with novel C-Terminal ending. **G.** Isoforms with N-Terminal isoforms. **H.** Isoforms with the novel exon 6b. **I.** PGC-1α-trE6-Ex13c **J.** sNT-PGC-1α **K.** PGC-1α-E3c **L.** CT-PGC-1α-E3c. The recovery of expression in the infarct zone after 16 days (ratio between 3d and 16d) under High-Fat-Diet is impaired, leading to a continuing downregulation of exon 1a and b. For the group-wise detectable isoforms, the downregulation is dominant in the infarct zone and less in the remote area in direct comparison of expression values. Most of the isoforms recover expression values compared over time with exception of the N-Terminal Isoforms in infarct area. Data acquired by qPCR using primers covering specific exon-exon-junctions. Expression values are normalized to housekeeper NUDC. n=4 each, Bars depict mean values, error bars represent SD. Significance calculated by unpaired student’s t-test (ns p≤0,05, *p≤0,05, **p≤0,01, ***p≤0,001).

**Table S01:**
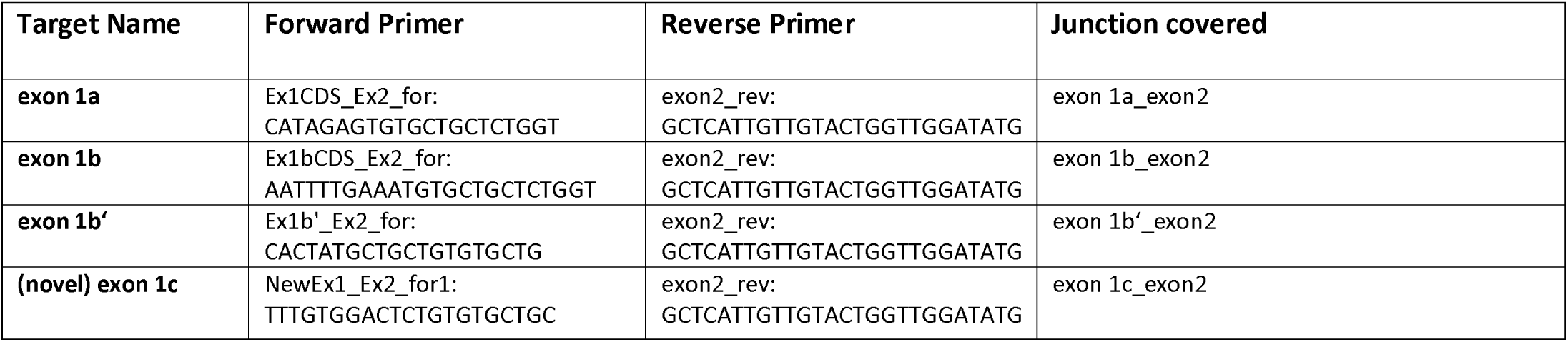
Primers used for detecting starting exons with q(PCR)

**Table S02:**
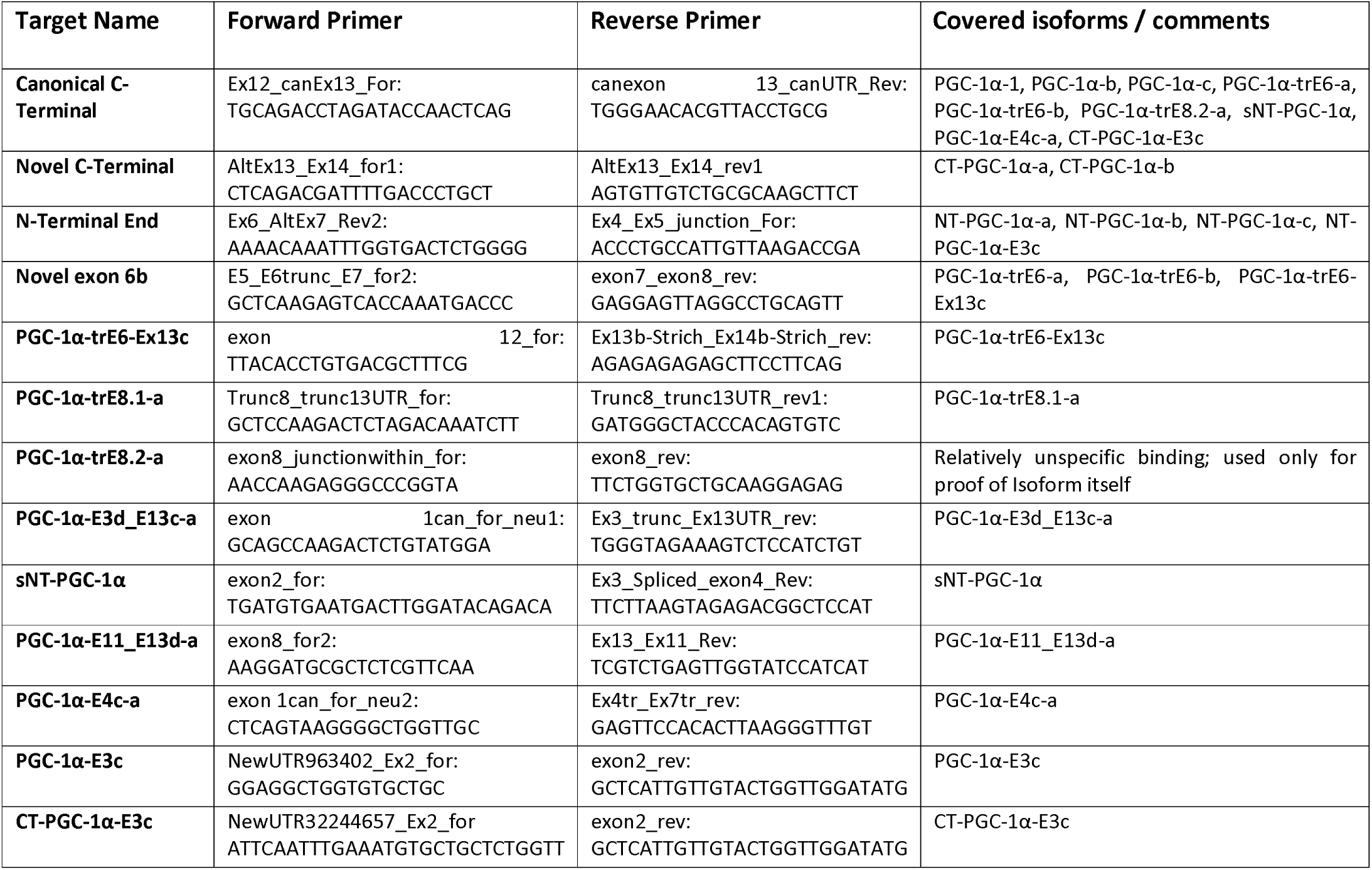
Primers used for detecting isoform-pattern with q(PCR)

## Supplemental Methods

### Ischemia/reperfusion injury (I/R) and sham-surgery in mice

In brief, mice were anesthetized with 2% isoflurane, orotracheally intubated, and ventilated with oxygen-enriched gas (40% oxygen) using a rodent ventilator (Minivent microventilator, Hugo Sachs, Germany). Mice were placed in a supine position on a warming plate (Uno, Zevenaar, the Netherlands) to keep body temperature at 37.5°C and received buprenorphine (0.1 mg/kg body weight, subcutaneously [s.c.]) for analgesia. Electrocardiography (ECG) was recorded continuously. After lateral thoracotomy, the pericardium was dissected, and a 7-0 surgical prolene suture was cautiously passed underneath the LAD coronary artery at a position 1 mm from the tip of the left auricle. The suture ends were passed through silicon tubing to form a snare occluder. Myocardial ischemia was produced by tightening the snare and confirmed by blanching of the myocardium and change in ECG (decrease in S wave amplitude). After 45 min, the snare occluder was opened to initiate reperfusion. The sham-operated controls underwent the same procedure but without ligation of the LAD. Afterward, the suture was removed, and the chest was closed. At the end of the experimental procedures, mice were extubated after they regained spontaneous breathing. Animals received buprenorphine (0.05–0.1 mg/kg body weight, s.c.) every 8 hr for up to 5 days for postoperative analgesia. Mice were excluded from the experiment when certain criteria of suffering were observed. These included weight losses greater than 20 % of body weight, cessation of food and water ingestion or lack of voluntary movement.

### cDNA Synthesis and quantitative RT-PCR

Shortly, the homogenate of tissue and RLT buffer was treated with proteinase K for 20 min before RNA was extracted on Qiagen RNeasy columns. A DNAse digestion took place on-column. Quality of eluted RNA was assessed by microvolume spectrophotometer analysis (Nanodrop™, Thermofisher Scientific). cDNA was synthesized from 1 μg RNA using the QuantiTect reverse transcription kit (QIAGEN).

### Glucose and insulin tolerance tests

For GTT, mice were fasted for 17 h, challenged with i.p. injected glucose (2 mg/g body weight) and blood glucose levels measured before and 15, 30, 45, 60 and 120 min after glucose injection using a blood glucose meter (Ascensia Contour, Bayer). For ITT, mice were fasted for 5 hours before i.p. injection of insulin (0.75 U insulin/kg body weight) and blood glucose levels measured before and 15, 30, 45, 60 and 90 min after insulin injection.

### Serum insulin ELISA

Blood was isolated by cardiac puncture, centrifuged at 2000x g for 20 min and respective serum analysed for insulin levels using the Ultra Sensitive Rat Insulin ELISA Kit (Crystal Chemicals) according to manufacturer’s protocol.

### Bioinformatic Pipeline for SMRT-Analysis

For primary analysis, the Isoseq3-Pipeline from Pacific Biosciences has been used; followed by a secondary Analysis Pipeline with open-source tools and modified scripts. This included mapping to genome with Minimap2(1), collapsing transcripts with cDNA Cupcake and Annotation with SQANTI2(2). Due to the nature of the algorithm behind, transcripts with very low abundance and therefore low count number will be excluded by the pipeline automatically within the polishing steps. Usually, if not focusing on a specific gene locus, it is impossible to detect whether how many in fact true transcripts get lost by that manner. Hence, interested in specifically PGC-1α-Isoforms, we created a second search strategy (Figure 1) by a similarity search strategy against sequence structures of the (previously) longest known transcript PGC-1α-a (= PGC-1α-1). Finding a balance between on one hand losing transcripts by clustering, merging and polishing transcripts and on the other hand analysing artificial, false-positive reads resulting from too low quality, we decided to use sequences from an intermediate step, called Full-Length-Non-Concatemere (FLNC) reads. Furthermore, starting from the raw data, only necessary quality control steps were performed to generate first circular-consensus reads (CCS) with primer removal and correct orientation (‘Full-Length” or ‘FL”-Reads) followed by Refinement (Poly-A-Trimming, Concatemer removal). Those FLNC were then used as database for the similarity search resulting in about 300 Sequences which were then manually annotated to known sequences of PGC-1α-transcripts using SnapGene(3).

### Motif annotation and prediction

For the Motif annotation and prediction, the following tools were used: NCBI’s Conserved Domain Database (CDD) (4), SMART protein domain annotation resource (5) and InterPro (6) as well as literature search on specific domain annotations of PGC-1α and detailed information on nuclear receptor domains (7–11). The promoter prediction was done using ElemeNT (12).

Analysis of the longest canonical isoform PGC-1α-a by literature search as well as motif annotation and prediction tools(4,6–11,13) exhibited existence of mainly involvement of six protein domains: A transcription activation domain (AD1(14), residues 30-40) accompanied by a second (AD2(14), residues 82-95), 3) a *LLXXLL*-motif(8,14,15) (residues 141–147) involved in transcriptional regulatory processes, a binding domain (residues 292–338) for interaction with the upstream target PPARγ (16), a RNA-Binding-motif(4,6,13) (residues 677–746), possibly involved in splicing processes of downstream mRNA targets and the PDB domain 3D24|D(8, 10) (residues 198-218), involved in binding of the estrogen-related receptor-alpha (ERRalpha).

### Scripts for SMRT Analysis

The following scripts have been used for analysis of SMRT-Data on the High-Performance Cluster using BioConda-Environment. As coding editor, “Sublime Text” for Windows (Version Build 4113) was used. The Scripts here containing the essential information on settings, please be aware that not every step (e.g. copying or moving of files) is mentioned.

#### 1. Creation of Circular Consensus Reads

Used CCS Version: 6.0.0 (commit v6.0.0-2-gf165cc26)

Using libraries:

unanimity : 6.0.0 (commit v6.0.0-2-gf165cc26)

pbbam : 1.6.1 (commit SEQII-release-10.0.0-35-g8fc7d89)

pbcopper : 1.8.0 (commit SEQII-release-10.0.0-38-g56f07ff)

boost : 1.73

htslib : 1.10.2

zlib : 1.2.11

Commandline:

~~~
ccs --log-level INFO -j 20 --log-file CCS_"$PROJECTNAME"_"$i"_logfile.txt --reportFile
CCS_"$PROJECTNAME"_"$i"_report.txt "$rawfiles" "$PROJECTNAME"_"$i".ccs.bam;
~~~

#### 2. Primer removal using LIMA-Tool

Used LIMA Version: lima 2.0.0 (commit v2.0.0)

Commandline:

~~~
lima "$PROJECTNAME"_"$i".ccs.bam "$primerpath" "$PROJECTNAME"_"$i"_demux.bam --isoseq --
peek-guess --dump-removed --dump-clips --log-level INFO --log-file
Lima_logfile_"$PROJECTNAME"_"$i".txt;
~~~

#### 3. Refine Step (Creation of FLNC reads)

Used Isoseq3-Version: isoseq3 3.4.0 (commit v3.4.0)

Commandline:

~~~
isoseq3 refine "$PROJECTNAME"_"$i"_demux.5p--3p.bam "$primerpath"
"$PROJECTNAME"_"$i"_flnc.bam --require-polya --log-level INFO --log-file
Refine_logfile_"$PROJECTNAME"_"$i".txt;
~~~

#### 4. Clustering Step

Used Isoseq3-Version: isoseq3 3.4.0 (commit v3.4.0)

Commandline:

~~~
isoseq3 cluster flnc.fofn "$PROJECTNAME"_flnc_clustered.bam --verbose --use-qvs --log-
level INFO --log-file Cluster_logfile_"$PROJECTNAME".txt
~~~

#### 5. Mapping of Long-Reads to Genome using minimap2

Used minimap2-Version: 2.18-r1015

Used Genome and annotation files:

##### Mice

- GRCm39.primary_assembly.genome.fa.gz (ENCODE)
- gencode.vM27.annotation.gtf.gz (ENCODE)

Note: For comparison reasons, analysis was also done additionally using ENSEMBL-Genome (Mus_musculus.GRCm39.dna.primary_assembly.fa.gz and Mus_musculus.GRCm39.104.chr.gtf.gz)

##### Human

- GRCh38.primary_assembly.genome.fa.gz (GENCODE)
- gencode.v38.annotation.gtf.gz (GENCODE)

Note: For comparison reasons, analysis was also done additionally using ENSEMBL-Genome (Homo_sapiens.GRCh38.dna.primary_assembly.fa.gz and Homo_sapiens.GRCh38.104.chr.gtf.gz)

Commandline:

~~~
minimap2 -t 20 -ax splice:hq -uf --secondary=no "$genomepath"/"$ensemblgene" \
"$PROJECTNAME"_flnc_clustered.fq > aligned_ensembl_"$PROJECTNAME"_flnc_clustered.sam 2> \
aligned_ensembl_"$PROJECTNAME"_flnc_clustered.sam.log
~~~

followed by sorting:

~~~
sort -k 3,3 -k 4,4n $DATEINAME.sam > sorted"$DATEINAME".sam;
~~~

#### 6. Collapsing isoforms using Cupcake-Tool

Used collapse_isoforms_by_sam.py from Cupcake-Tools and seqkit.

Commandline:

Removing Duplicates:

~~~
seqkit rmdup -D duplicate_list_"$DATEINAME".txt "$DATEINAME".fq > "$DATEINAME"_uni.fq;
~~~

B) Collapse Transcripts:

~~~
collapse_isoforms_by_sam.py --input "$DATEINAME".fq --fq -s "$p" --dun-merge-5-shorter -o "$p";
~~~

#### 7. SQANTI3 for Reporting of Isoforms and Gene Usage

Used R scripting front-end version 3.6.1 (2019-07-05) as well as SQANTI Version 3.0

Commandline:

~~~
sqanti3_qc.py --gtf "$p" "$PROJECTNAME"/SQANTI3/genome_SQ_G/"$gencodeannotation_SQ"
"$PROJECTNAME"/SQANTI3/genome_SQ_G/"$gencodegene_SQ" --aligner_choice=minimap2 -t 20;
~~~

